# Two bi-functional cytochrome P450 CYP72 enzymes from olive (*Olea europaea*) catalyze the oxidative C-C bond cleavage in the biosynthesis of secoxy-iridoids - flavor and quality determinants in olive oil

**DOI:** 10.1101/2020.08.28.265934

**Authors:** Carlos E. Rodríguez-López, Benke Hong, Christian Paetz, Yoko Nakamura, Konstantinos Koudounas, Valentina Passeri, Luciana Baldoni, Fiammetta Alagna, Ornella Calderini, Sarah E. O’Connor

**Author notes:** To whom correspondence should be addressed (SOC); (OC), (FA).

## Abstract

- Olive (*Olea europaea*) is an important crop in Europe, with high cultural, economic, and nutritional significance. Olive oil flavor and quality depend on phenolic secoiridoids, but the biosynthetic pathway of these iridoids remains largely uncharacterized.
- We discovered two novel, bi-functional cytochrome P450 enzymes, catalysing the rare oxidative C-C bond cleavage of 7-epi-loganin to produce oleoside methyl ester (OeOMES) and secoxyloganin (OeSXS), both through a ketologanin intermediary. Although these enzymes are homologous to the previously reported *Catharanthus roseus* Secologanin Synthase (CrSLS), the substrate and product profiles differ.
- Biochemical assays provided mechanistic insights into the two-step OeOMES and CrSLS reactions. Model-guided mutations of OeOMES changed the product profile in a predictable manner, revealing insights into the molecular basis for this change in product specificity.
- Our results suggest that, in contrast to published hypotheses, *in planta* production of secoxy-iridoids is secologanin independent. Notably, sequence data of cultivated and wild olives, points to a relation between domestication and OeOMES expression. Thus, the discovery of this key biosynthetic gene suggests a link between domestication and secondary metabolism, and could potentially be used as a genetic marker to guide next-generation breeding programs.

## Introduction

Olive (*Olea europaea* L.) is an economically and culturally important crop in Europe. The European Union produces two thirds of the global supply of olive oil, and also consumes half of it (European Commission, 2020). Olives, along with wine and bread, are an important part of the Mediterranean diet, in the form of both its fermented, pickled fruit and its oil, which has been consumed for millennia. The specialized metabolites of olive fruit and oil have been associated with various health benefits (George *et al*., 2019; Schwingshackl *et al*., 2019; de la Torre *et al*., 2020; Marrero *et al*., 2020). The secoiridoids are the major class of metabolites present in olive fruit, with oleuropein, a phenolic secoiridoid, making up 6-14% of the dry weight of the fruit (Amiot *et al*., 1986; Ryan *et al*., 1999). Oleuropein strongly influences the quality of olive products because of its bitterness, which is a desirable trait in olive oil, though table olives must be subjected to a lengthy curing process to degrade this compound (Garrido-Fernández *et al*., 1997). In planta, the glucose of oleuropein can be hydrolyzed by a dedicated glucosidase (Koudounas *et al*., 2015, 2017; Velázquez-Palmero *et al*., 2017), as part of a defence response. The resulting aglycone has protein-crosslinking and lysine-alkylating activities, making this compound an effective deterrent of generalist herbivores (Konno *et al*., 1999).

Obtaining olive varieties with “fine-tuned” oleuropein levels is an important goal of olive breeding programs (Pérez *et al*., 2018). The benefits of secoiridoid-related markers have been recently highlighted in the industrial processing of table olives, where oleuropein degradation via glucosidases was a major biochemical marker for strain selection (Bavaro *et al*., 2017). The importance of secoiridoids in systematic breeding of olive for quality Virgin Olive Oil is increasingly being recognized, with lack of knowledge on their genetic determinants as the major hurdle for this cause (Pérez *et al*., 2018). Since genotype appears to be a stronger determinant of secoiridoid contents than environmental conditions (Pérez *et al*., 2018), identification of the genes responsible for oleuropein biosynthesis could readily benefit the olive oil industry as a molecular marker.

Despite the agricultural, biomedical and ecological importance of iridoids, the biosynthetic pathway of only two iridoids has been solved to date: nepetalactone, an evolutionarily and structurally unrelated iridoid produced in catnip (Lichman *et al*., 2019; Lichman *et al*., 2020), and secologanin, a secoiridoid produced by *Catharanthus roseus* and several plants in the Gentianales order (Irmler *et al*., 2000; Salim *et al*., 2013; Miettinen *et al*., 2014). Secoiridoids are characterized by lack of the cyclopentane ring, which is subjected to an oxidative cleavage reaction. This reaction is catalyzed by the enzyme secologanin synthase (SLS), a cytochrome P450 (CYP72) first identified from *C. roseus*, which catalyzes the oxidative opening of loganin into secologanin (Irmler *et al*., 2000). The only additional homolog that has been characterized is secologanic acid synthase from *Camptotheca acuminata*, which catalyzes the same oxidative cleavage as SLS, but using des-methylated loganin (loganic acid) as the substrate (Yang *et al*., 2019). Unlike secologanin-type iridoids, in which the oxidative cleavage produces a vinyl group, oleoside-type iridoids have a distinctive exocyclic olefin (Fig. 1), which increases the reactivity of the aglycone for defensive protein crosslinking (Konno *et al*., 1999). Oleoside-type iridoids are a chemotaxonomic marker, being produced exclusively in the Oleaceae family, with olive producing both secologanin- and oleoside-type secoxy-iridoids.

**Fig. 1.**
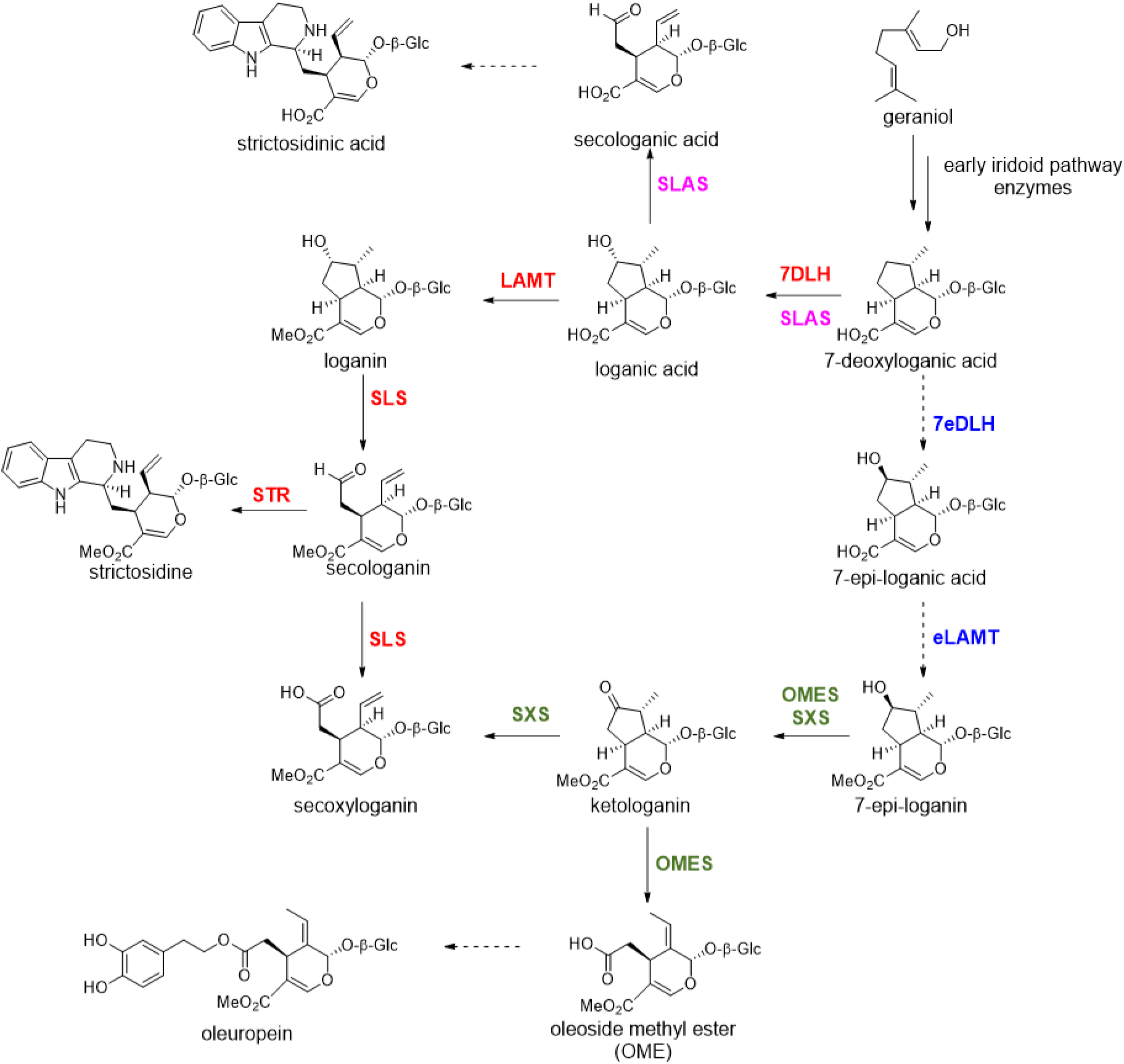
Summary of existing knowledge of the secoiridoid pathways. Enzymes in red have been characterized in *Catharanthus roseus*, enzymes in pink have been characterized in *Camptotheca acuminata*. Enzymes in blue are hypothesized to exist in *Olea europaea*, but not in *C. roseus*. Dashed arrows indicate steps for which a biosynthetic gene has not yet been discovered. Enzymes in olive green were characterized in this work in *O. europaea*. SLS, secologanin synthase; SLAS, secologanic acid synthase; STR, strictosidine synthase; 7DLH, 7-deoxy-loganic acid hydroxylase; LAMT, loganic acid methyl-transferase; 7eDLH, 7-epi-deoxy-loganic acid hydroxylase; eLAMT, 7-epi-loganic acid methyl-transferase; OMES, oleoside methyl ester synthase; SXS, secoxyloganin synthase; OS, oleuropein synthase.

The structure of oleuropein and secologanin suggests that the early biosynthetic steps of these two compounds are the same; namely biosynthesis of geraniol, cyclization to nepetalactol, followed by oxidation and glucosylation (Fig. 1). The identification of olive Iridoid Synthase (OeISY) supports this hypothesis (Alagna *et al*., 2016). However, the mechanism of the chemotaxonomical divergence of secologanin- and oleoside-type iridoids remains unknown. Several biosynthetic precursors for oleuropein have been proposed: secologanin (Inouye *et al*., 1971; Damtoft *et al*., 1993; Alagna *et al*., 2016), 8-epi-kingiside (Inouye *et al*., 1971; Damtoft *et al*., 1993), or ketologanin (Alagna *et al*., 2016) (Fig. S1). Although several feeding experiments to olive plants have been performed (Inouye *et al*., 1971; Damtoft *et al*., 1993; Serrilli *et al*., 2016), no conclusive evidence for the oleoside pathway has been established.

Here, we use olive transcriptomic data to identify the enzymes responsible for generating the secoiridoid scaffolds in olive. Using SLS from *C. roseus* as a bait for a homology based search, we identified three SLS homologs in olive, and showed that these SLS homologues either catalyze formation of Oleoside Methyl Ester (OeOMES) or secoxyloganin (OeSXS) from 7-epi-loganin, through a ketologanin intermediary that can also be directly uptaken by the enzymes. This work reports the key step for this important flavor agent and mutational analysis reveals how enzyme substrate and product specificity are modulated to generate chemical diversity in plants.

## Materials and Methods

### Identification of enzyme candidates

To identify candidates for oleoside methyl ester biosynthesis, we blasted the protein sequence of *Catharanthus roseus* Secologanin Synthase 2 (CrSLS2) against a previously published, in house assembled Olive transcriptome (Alagna *et al*., 2009), from which expression correlations were also calculated. We designed primers for the three homologous sequences found (*OeSLS1-3*; GenBank MT909126, MT909123, and MT909125, respectively) and amplified them via PCR from a cDNA library derived from total RNA of Olive (*O. europaea* subs. *europaea*) young leaf tissue. Since we were unable to obtain full-length clones of *OeSLS1* and *OeSLS3*, DNA fragments were ordered from IDT (Integrated DNA Technologies, BVBA; Belgium). The sequences were cloned into pUC19 and sequence verified, and later sub-cloned using In-Fusion cloning system (Takara Bio Europe) into the Gal10 MCS of pESC-Leu2d containing an *Artemisia annua* Cytochrome P450 reductase (AaCPR) under the Gal1 promoter (Ro *et al*., 2008). Mutants were directly synthesized in the pESC-Leu2d::AaCPR by Twist (Twist Bioscience, USA).

### Heterologous expression and microsomal preparations

Protease deficient *Saccharomyces cerevisiae* strain YPL 154C:Pep4KO was used for heterologous expression of p450s, as previously published (Ro *et al*., 2002), with minor modifications. Namely, the strain, harboring the dual expression pESC-Leu2d::AaCPR-GOI plasmid, was plated, and single colonies inoculated in 5 mL Synthetic Defined medium without leucine (SD-Leu; Sigma-Aldrich) with 2% glucose, and cultivated overnight in an orbital shaker at 30°C and 250 rpm. This culture was used to inoculate 100mL of SD-Leu with 2% glucose, left to grow for 24 hours. This culture was diluted into 1 L SD-Leu with 2% galactose, for protein expression, to a final absorbance of OD_600_ = 0.5 (~6-7×10^6^ cells/mL). After 24 hours, cells were harvested, and microsomes prepared as previously mentioned (Dang *et al*., 2017), using an LM20 microfluidizer (Siemens).

Microsome reactions were performed using 2 mg of total protein in 50 μL, in 50 mM Tris buffer at pH 7.5, incubated at 30°C in the dark on a shaker (Vortemp56, Labnet) at 300 rpm. Exploratory reactions were incubated overnight with 25 μM substrate and 125 μM NADPH. Steady-state kinetics were determined by feeding 10, 20, 40, 80, 160, 250, and 500 μM substrate, with thrice as much NADPH, and stopping the reactions at 30, 60, 90, 120 and 180 minutes. All reactions were done in triplicate, started by adding the substrate, and stopped by adding 50 μL methanol. Samples were then centrifuged at 20,000 RCF for 10 minutes and transferred to HPLC vials or plates.

### Iridoid profiling using HPLC-MS

Samples were analyzed using an Elute LC system (Bruker Daltonik, Bremen, Germany) coupled via ESI to a Maxis II q-TOF (Bruker Daltonik). Iridoids were separated using an Acquity UPLC^®^ C18 column (2.1×50 mm, 1.7 μm, 100 Å; Waters) at 40 °C with a gradient from water (A) to acetonitrile (B), both modified with 0.1% formic acid, using two different programs, both at a flow of 0.3 mL/min and redirecting to waste the first 90 seconds. For exploration of reaction products, an 11 minute gradient was used, starting at 1% B for one minute, then linearly increasing to 20% B in five minutes, then to 40% B in two minutes, and finally to 99% B in one minute, and kept for two minutes at 99% B. The column was equilibrated in 1% B for another two minutes before the next injection. For the kinetic experiments, the gradient was shortened to 4.5 minutes, starting at 1% B for 0.5 minutes, then linearly increasing to 20% B in 2.5 minutes, then 25% B in 0.5 minutes, and immediately changing to 99% B, which was kept for one minute. The column was equilibrated in 1% B for one minute.

Acquisition of MS data was performed using the same conditions for exploration and kinetics, but the latter with no fragmentation. Ionization was performed via pneumatic assisted Electro Spray Ionization in negative mode (ESI-) with a capillary voltage of 3.5 kV, an end plate offset of 500 V, and a nebulizer pressure of 3 bar. Nitrogen gas was used as frying gas at 350°C and a flow of 11 L/min. For the kinetics experiments, acquisition was performed at 2 Hz with no fragmentation, at a mass range from 250 to 1000 m/z. For exploration experiments, acquisition was done at 12 Hz following a mass range from 250 to 1000 m/z, with data dependent MS/MS without active exclusion. Fragmentation was triggered on an absolute threshold of 400, and acquired on the most intense peaks using a target intensity of 2×10^4^ counts, with a MS/MS spectra acquisition of 12 Hz, and limited to a total cycle time range of 0.5 s, and a dynamic collision energy from 20 to 50eV. At the beginning of each run, while the LC input was redirected to waste, a sodium formate-isopropanol solution was injected at 18 mL/h for the first 90 seconds of each run, and the m/z values were re-calibrated using the expected cluster ion m/z values.

### Data analysis

Raw MS files were converted to mzXML using Bruker Data Analysis software (Bruker Daltonik, Bremen, Germany) and, when needed, Extracted Ion Chromatograms (XIC) were exported to Excel^®^ using MZMine2 (v2.40.1; Pluskal *et al*., 2010). For the kinetics experiment, areas were obtained by an in-house built script, using the R programming language (R Core Team, 2019). The script accessed the raw data files, extracted the XIC within 0.05 m/z error, and integrated it in a window of 0.1 minutes within the retention time of each standard. Raw mzXML files were accessed with the aid of the *mzR* package (Chambers *et al*., 2012) and parallel processing of data facilitated by the *parallel* (R Core Team, 2019) and *snow* (Tierney *et al*., 2018) libraries. Figures were done using the R base package (R Core Team, 2019), swarmed boxplots using the *beeswarm* library (Eklund, 2016) and chromatograms using Excel^®^. Molecules were drawn using ChemDraw^®^.

### Steady-State Kinetics

For quantification in the kinetics experiments, a ten-point calibration curve of 7-epiloganin, ketologanin, and oleoside methyl ester, from 0.1 to 100 p mol, was run in between each replicate of all substrate concentrations for each enzyme. Concentrations were quantified with the enclosing calibration curves of each sample; secoxyloganin was quantified in OME equivalents. Linear velocities were calculated by running linear regressions, and steady-state constants were estimated by a non-linear least squares fit to a Michaelis-Menten equation or Eq. 1, according to the case, using the R *stats* library (R Core Team, 2019). Derivation of Eq. 1 is shown in Notes S1 and S2.

### Homology models

Protein sequences of biochemically tested CYP72 enzymes were obtained from Uniprot (The Uniprot Consortium, 2019), multiple alignments were performed using ClustalW (Madeira *et al*., 2019), and phylogenetic trees generated using the IQ-TREE web server (Trifinopoulos *et al*., 2016) and visualized using the interactive Tree of Life (Letunic & Bork, 2019). The homology model for OeOMES, OeSXS, CrSLS and CAC CYP72A612 was selected using PHYRE2 online software (Kelley *et al*., 2015) and all models were aligned using POSA (Li *et al*., 2014). For visual exploration, and selection of the regions closest to the substrate, we used UCSF Chimera v1.14 (Pettersen *et al*., 2004).

### Analysis of RNA-Seq public datasets

We selected publicly available RNA-Seq data by searching in NCBI’s Sequence Read Archive, and downloaded the raw fastq files of projects PRJNA525000 (Gros-Balthazard *et al*., 2019), PRJNA256033 (Leyva-Pérez *et al*., 2015), PRJNA378602 (Jiménez-Ruiz *et al*., 2019), PRJNA401310 (Jiménez-Ruiz *et al*., 2018), PRJNA514943 (Research Institute of Forestry), and PRJNA596876 (Research Institute of Forestry) using the European Nucleotide Archive FTP. Salmon (Patro *et al*., 2017) was used for abundance estimation via quasi-mapping of reads against transcriptomes derived from the olive Farga cultivar genome (Cruz *et al*., 2016), wild olive genome (Unver *et al*., 2017), and the Reprolive project (Carmona *et al*., 2015), generated from vegetative and reproductive tissue. The Farga cultivar transcriptome was chosen for analysis, as it was the only one with mappings of more than 70% (Fig. S13). Of the 112 samples, 103 had a mapping greater than 70 %, having a total of 2×10^9^ mapped reads, an 82% mapping total (Fig. S13).

### Iridoid standards

Loganin and secologanin standards were purchased from Sigma-Aldrich, and secoxyloganin was obtained by incubating secologanin and CrSLS microsomes overnight, with five NADPH equivalents. 8-epiloganin as well as oleoside methyl ester were purchased from AnalytiCon Discovery GmbH. An inconsistency exists in the PubChem database and AnalytiCon Discovery catalogue, both reporting oleoside methyl esther with the exocyclic bond in the Z configuration, in contrast to the published literature (Inouye *et al*., 1971; Damtoft *et al*., 1993; Alagna *et al*., 2016) which report the E configuration. Thus, the determination of the absolute configuration of the oleoside methyl esther standard was performed by NMR (Methods S1), which confirmed that the standard from AnalytiCon Discovery is in the E configuration (Fig S14 and S15), despite it being reported in the Z configuration in their catalogue.

The rest of the iridoid standards are not commercially available, and were synthesized. The detailed synthesis protocols, along with yields and spectroscopic data, are shown in the supplementary methods (Methods S2). The spectroscopic data are in accordance with the literature values reported for geniposide pentaacetate (Zhang *et al*., 2013); 10-deoxy-geniposide tetraacetate, 10-deoxygeniposidic acid, and 7-deoxy-loganic acid (Inoue *et al*., 1992); 7-deoxy-8-*epi*-loganic acid (Nakamura *et al*., 2000); loganin epoxide and ketologanin tetraacetate (Inouye *et al*., 1970); 7-keto-loganin (Gross *et al*., 1986); and 7-*epi*-loganin (Itoh *et al*., 2005). Spectroscopic data for 7-*epi*-8-*epi*-loganin, which has not been previously reported, is presented below and in Fig S16 and S17. 7-*epi*-8-*epi*-loganin: ^1^H NMR (400 MHz, CD_3_OD) δ 7.43 (s, 1H), 5.66 (d, *J* = 5.6 Hz, 1H), 4.70 (d, *J* = 7.9 Hz, 1H), 4.17 (dd, *J* = 12.7, 5.7 Hz, 1H), 3.89-3.84 (m, 1H), 3.69 (s, 3H), 3.68-3.61 (m, 1H), 3.39-3.34 (m, 1H), 3.24-3.17 (m, 3H), 2.84 (dd, *J* = 18.0, 8.4 Hz, 1H), 2.61 (ddd, *J* = 14.2, 9.8, 7.4 Hz, 1H), 2.22 (td, *J* = 12.1, 6.6 Hz, 2H), 1.42-1.32 (m, 1H), 1.10 (d, *J* = 7.0 Hz, 3H);

^13^C NMR (100 MHz, CD_3_OD) δ 168.2, 151.2, 112.2, 98.5, 95.7, 76.9, 76.6, 74.0, 73.5, 70.3, 61.5, 50.3, 42.2, 41.7, 40.5, 30.9, 9.4;

## Results

### Identification of enzyme candidates for secoiridoid formation in Olea europaea

We identified 3 SLS (*C. roseus*) homologs, identified as *OeSLS1-3*, in the olive transcriptome (Fig. 2). A co-expression analysis using transcriptome data from different olive tissues showed that, while *OeSLS1* showed modest correlation (Pearson’s r=0.48), *OeSLS2* and *OeSLS3* showed high co-expression with the olive Iridoid Synthase (r=0.76 and r=0.95, respectively). OeSLS1-3 were cloned, heterologously expressed in yeast and the subsequent yeast microsomes were assayed for biochemical activity. During amplification of these genes from olive cDNA, it became evident that *OeSLS2* had a variant, *OeSLS2A*, with 99.8% identity to *OeSLS2*. Subsequent enzyme assays indicated that OeSLS2A had identical biochemical activity as OeSLS2, and was not investigated further.

**Fig. 2.**
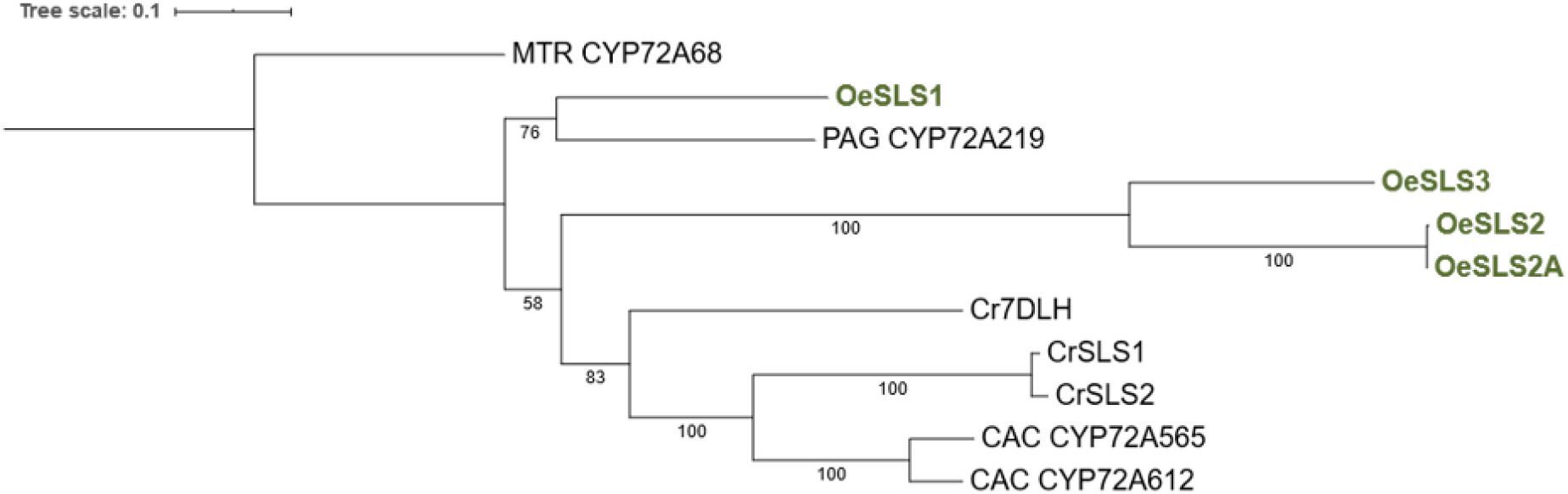
Consensus alignment tree. Maximum-likelihood tree (LG+G4) of a ClustalW alignment of the protein sequences of CYP72 enzymes with proven biochemical activity. Nodal support was estimated using a 1000 bootstrap, and the tree rooted on MTR CYP72A68. *Catharanthus roseus* (Cr) SLS and *Camptotheca acuminata* (CAC) CYP72 enzymes catalyze oxidative cleavage of iridoid scaffolds to produce secologanin and secologanic acid, respectively, while *Panax ginseng* (PAG) and *Medicago truncatula* (MTR) CYP72 enzymes oxidize triterpenoid scaffolds, without cleavage. The enzymes covered in this work for *Olea europaea* (Oe) are shown in bold olive green.

### Biochemical functions of SLS homologues

Oxidative cleavage of the iridoid scaffold is required to generate the secoiridoids. Feeding studies with olive plants (Inouye *et al*., 1971) indicate that the final product of such an oxidative cleavage product would be oleoside methyl ester, which is then esterified to form oleuropein. The most straightforward hypothesis is that a secologanin synthase (SLS) homologue would catalyze such a reaction from an iridoid intermediate, but no experimental data suggested what specific substrate this putative enzyme would utilize as a starting material. The two characterized oxidative cleavage reactions in secoiridoid biosynthesis, namely SLS from *C. roseus* and SLAS from *C. acuminata*, utilize loganin and loganic acid, respectively (Fig. 1). Several substrates have been suggested for oleoside methyl ester (OME) biosynthesis, such as secologanin (Inouye *et al*., 1971; Damtoft *et al*., 1993; Alagna *et al*., 2016), 8-epi-kingiside (Inouye *et al*., 1971; Damtoft *et al*., 1993), and ketologanin (Alagna *et al*., 2016) (Fig. S1).

We first assayed the olive SLS homologues with secologanin and loganin (Fig. S2). SLS from *C. roseus* (CrSLS) was used for comparison. While OeSLS1 did not show activity, CrSLS, OeSLS2 and OeSLS3 converted secologanin to secoxyloganin, although the latter to a less extent (Fig. S2a). Notably, all three SLS homologues (OeSLS1, OeSLS2, and OeSLS3) failed to react with loganin (Fig. S2b), the native substrate of CrSLS. Interestingly, although secoxyloganin production has been reported for CrSLS (de Bernonville *et al*., 2015), *C. roseus* has no detectable concentrations of secoxyloganin; in contrast, secoxyloganin is abundant in olive (De Marino, *et al*., 2014).

Since these enzymes failed to produce oleoside methyl ester from loganin and secologanin, we next assayed SLS homologues with ketologanin. Ketologanin was not implicated in any known secoiridoid biosynthetic pathway, but previous research (Damtoft *et al*., 1993; Alagna *et al*., 2016) has suggested that this compound may be a biosynthetic intermediate in oleoside biosynthesis. While CrSLS and OeSLS1 failed to react with this substrate, both OeSLS2 and OeSLS3 turned over ketologanin; OeSLS2 oxidatively cleaved ketologanin to oleoside methyl ester (OME) and OeSLS3 oxidatively cleaved it to secoxyloganin (Fig. 3a). We thus renamed OeSLS2 Oleoside Methyl Ester Synthase (OeOMES) and OeSLS3 Secoxyloganin Synthase (OeSXS).

**Fig. 3.**
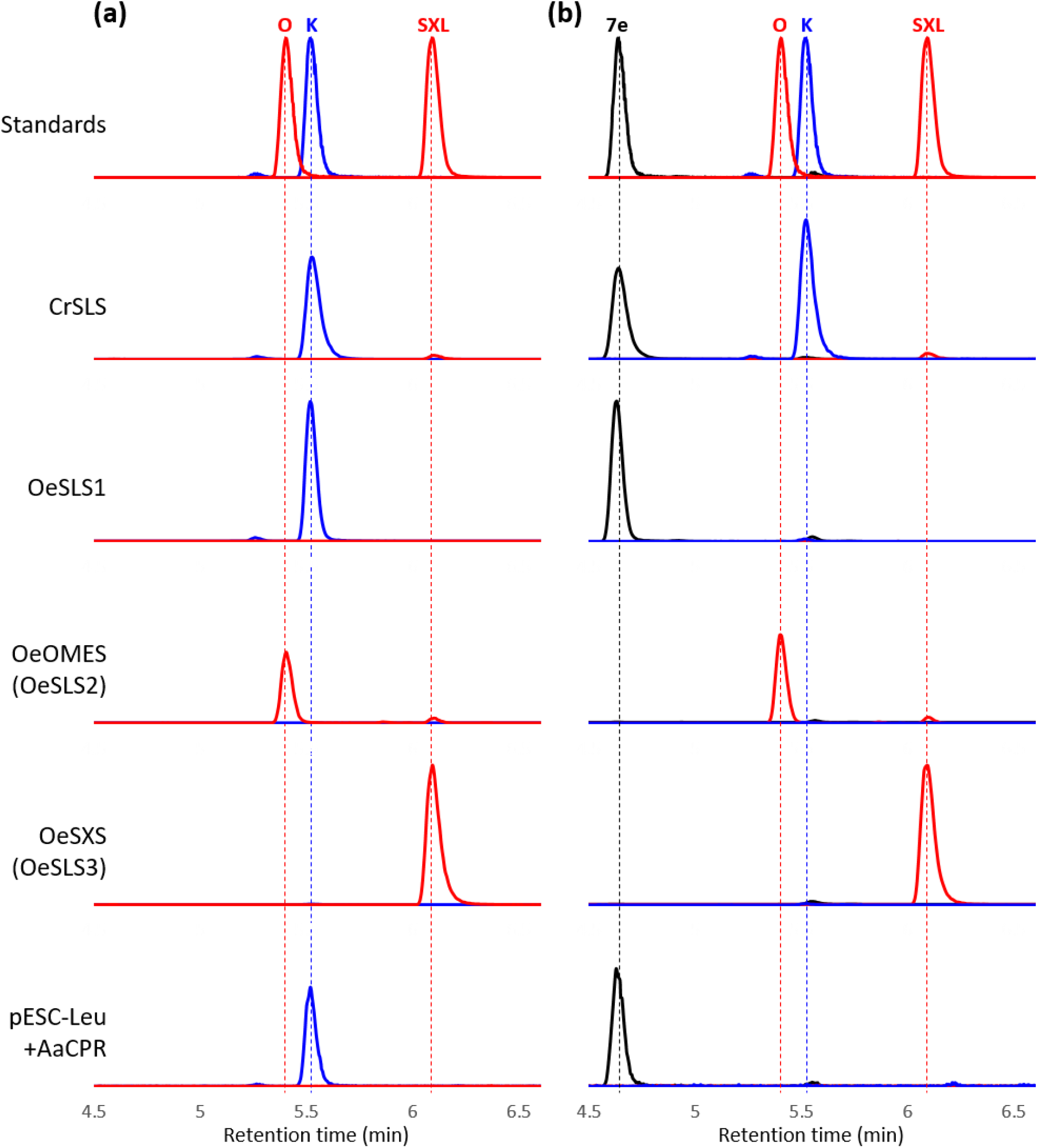
Extracted Ion Chromatograms (XIC) of microsome substrate assays. XIC of the microsomal incubations with ketologanin (a) and 7-epi-loganin (b) are shown. Three channels are depicted, corresponding to the most abundant adduct of OME or secoxyloganin (red, [MH]^−^ = 403.1240±0.05), ketologanin (blue, [M+FA-H]^−^ = 433.1346±0.05) and 7-epiloganin (black, [M+FA-H]^−^ = 435.1503±0.05). Intensities are scaled to the highest intensity of the corresponding channel in all incubations. pESC-Leu_AaCPR refers to microsomes that express only the cytochrome reductase. O, oleoside; K, ketologanin; SXL, secoxyloganin.

Given that CrSLS can perform the sequential oxidation of loganin to secologanin to secoxyloganin, we next investigated whether the olive SLS homologues were also capable of performing two sequential oxidation steps. We synthesized a mixture of loganin diastereomers (7-epi-loganin, 8-epi-loganin, and 7,8-epi-loganin) to test if they could be oxidized by the SLS homologues. Of these epimers, only 7-epi-loganin was depleted in OeOMES and OeSXS, where peaks corresponding to OME and secoxyloganin, respectively, were detected (Fig. S3). Ketologanin was also detected in OeOMES and OeSXS reactions, and also, interestingly, when CrSLS was reacted with 7-epi-loganin. In these enzymatic reactions, an isomer of ketologanin was observed along with the ketologanin product (Fig. S3), although this product could not be isolated in sufficient quantity for characterization. OeSLS1, as with the other substrates, was largely inactive, but did produce trace amounts of ketologanin. These enzyme assays were then repeated with purified 7-epi-loganin, which confirmed the observation that OeOMES and OeSXS produce OME and secoxyloganin, respectively, while CrSLS produces ketologanin (Fig. 3b). Although only trace amounts of ketologanin were detected in long incubations with 7-epi-loganin and OeOMES and OeSXS, its presence in the early time points of the kinetic reactions supports its role as an intermediary.

We next tested 7-deoxyloganin to assess whether these enzymes could catalyze the three step oxidation: 7-deoxyloganin to 7-epi-loganin to ketologanin, and then finally to either OME or SXS. When incubated with 7-deoxyloganin, CrSLS produced loganin, subsequently oxidizing it to secologanin and secoxyloganin (Fig. S4). Notably, *C. roseus* has a dedicated hydroxylase (7DLH) that converts 7-deoxyloganic acid to loganic acid that has been validated by gene silencing (Salim *et al*., 2013), so it is not clear whether CrSLS turns over 7-deoxyloganin *in planta*. OeOMES oxidized 7-deoxyloganin to a product that could not be isolated in sufficient quantity for characterization. However, since only traces of ketologanin and OME were observed in this reaction, we assume that 7-deoxyloganin is not on pathway (Fig. S4). OeSXS was also found to produce this molecule, and OeSLS1 showed no activity. No activity was observed when the microsomes were incubated with 7-deoxyloganic acid, 7-deoxy-8-epiloganic acid, and 7-deoxy-8-epi-loganin.

### Steady-state kinetics of OeOMES and OeSXS

Kinetic parameters were calculated for OeOMES and OeSXS, incubated with either ketologanin and 7-epiloganin (Fig. S5). If enzyme regeneration is faster than the oxidation reaction, conditions that should be fulfilled by adding excess NADPH, the kinetic parameters for the ketologanin substrate should follow Michaelis-Menten kinetics (Michaelis & Menten, 1913). The exponential fit obtained (Fig. S5a,c) suggests that this assumption is reasonable, and can be used as a starting point to measure the kinetics of the consecutive reaction from 7-epiloganin. The subject of the dissociation of intermediates in consecutive reactions catalyzed by cytochrome P450 enzymes remains contentious, but the kinetics of either case do not practically differ under a quasi-steady-state assumption (Notes S1 Eq.S1.7 and Notes S2 Eq. S2.13), and follow the equation:

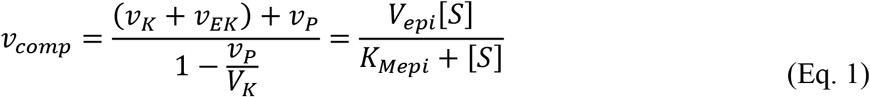

where [*S*] is the concentration of 7-epiloganin, *V_epi_* and *V_K_* are the maximum velocities of 7-epi-loganin and ketologanin, respectively; (*v_K_* + *v_EK_*) is the change in time of measured ketologanin (in solution plus complexed with the enzyme), and *v_P_* the rate of production of OME (by OeOMES) or secoxyloganin (by OeSXS). For simplification, the scaled sum of velocities is called *v_comp_* and follows Michaelis-Menten kinetics (Fig. S5b,d). The results of this model are shown in Fig. S5b for OeOMES and Fig. S5d for OeSXS. Individual velocities, used to calculate Fig. S5b,d, can be seen in Fig. S6 for OeOMES (Fig. S6a) and OeSXS (Fig. S6b).

The relative turnover 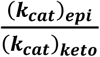 and specificity constants (V_max_/K_M_) of epiloganin versus ketologanin oxidation were compared (Table 1). These kinetic analyses indicate that OeOMES is more specific towards 7-epi-loganin and the V_max_ for oxidation of epiloganin to ketologanin is twice as fast than the oxidative cleavage to OME. This explains the non-Michaelis-Menten kinetics seen in the individual velocities (Fig. S6a), where as there is more substrate presence (>250uM epiloganin) the rate of ketologanin formation is higher than that of OME. In contrast, OeOSXS, although it has a similar relative specificity for both 7-epi-loganin and ketologanin, it has a lower relative turnover for 7-epi-loganin, indicating that the oxidative cleavage of ketologanin to secoxyloganin (V_max_ = 54.6±3.6 μM/s) is faster than the formation of ketologanin from 7-epiloganin (V_max_ = 12.6±0.6 μM/s). Again, this can be seen in the individual kinetic analyses where, unlike the OeOMES, the rate of secoxyloganin is always higher than the rate of ketologanin formation (Fig. S6b).

**Table 1.** Kinetic parameters of OeOMES and OeSXS. Values are shown as the calculated regression coefficient plus/minus the standard error. The relative turnover is the ratio of *k*_cat_ and the relative specificity is the specificity constant (*k*_cat_/K_M_) of epiloganin with respect to ketologanin.

| | 7-epi-loganin | | | Ketologanin | | | $\frac{(k_{cat})_{epi}}{(k_{cat})_{keto}}$ |
| --- | --- | --- | --- | --- | --- | --- | --- |
| | $K_M$ (uM) | $V_{max}$<br>(uM/s) | $V_{max}/K_M$<br>(s<sup>-1</sup>) | $K_M$<br>(uM) | $V_{max}$<br>(uM/s) | $V_{max}/K_M$<br>(s<sup>-1</sup>) | |
| **OeOMES** | 170±24 | 90.0±5.4 | 0.53±0.08 | 172±37 | 39.0±3.6 | 0.23±0.05 | 2.31±0.25 |
| **OeSXS** | 37.7±9.3 | 12.6±0.6 | 0.33±0.08 | 254±36 | 54.6±3.6 | 0.21±0.03 | 0.23±0.022 |

### Mutational analysis of OeOMES

Structural information for plant cytochrome P450s is limited, which hampers rational mutational analysis. Nevertheless, we decided to explore the effects of amino acid substitution, in an attempt to identify the molecular basis of substrate and/or product specificity. Since OeOMES can produce both secoxyloganin (from secologanin) and oleoside methyl ester (from ketologanin), we used OeOMES as a template to introduce amino acid mutations. Using Phyre2.0 (Kelley *et al*., 2015), we modelled OeOMES, OeSXS, CrSLS and CAC CYP72A612 against the crystal structure of *Saccharomyces cerevisiae* lanosterol 14-a demethylase (PDB 4LXJ; Monk *et al*., 2014), as it was the best compromise in similarity of all enzymes. This enzyme also catalyzes oxidative cleavage of a carbon-carbon bond, resulting in formation of a vinyl group and a carboxylic acid (Monk *et al*., 2014).

The resulting homology models were then aligned using POSA (Li *et al*., 2014), and we selected five regions of interest within 10Å of the substrate binding site or the iron reactive center (Fig. S7). As expected, the conserved region of CYP responsible of the coordination of iron aligned perfectly with the heme group in all the models. Within the 5 regions closest to the substrate and iron, named A-E, we chose the amino acids that were not conserved among OeOMES, OeSXS, CrSLS and CAC CYP72A612. We substituted all these amino acids in each region of OeOMES with the corresponding residues of OeSXS, yielding six mutant designs (Table S1). Each of these mutants was expressed in yeast, and the resulting microsomal fractions were incubated with ketologanin, 7-epiloganin and secologanin (Fig. 4; Fig. S8).

**Fig. 4.**
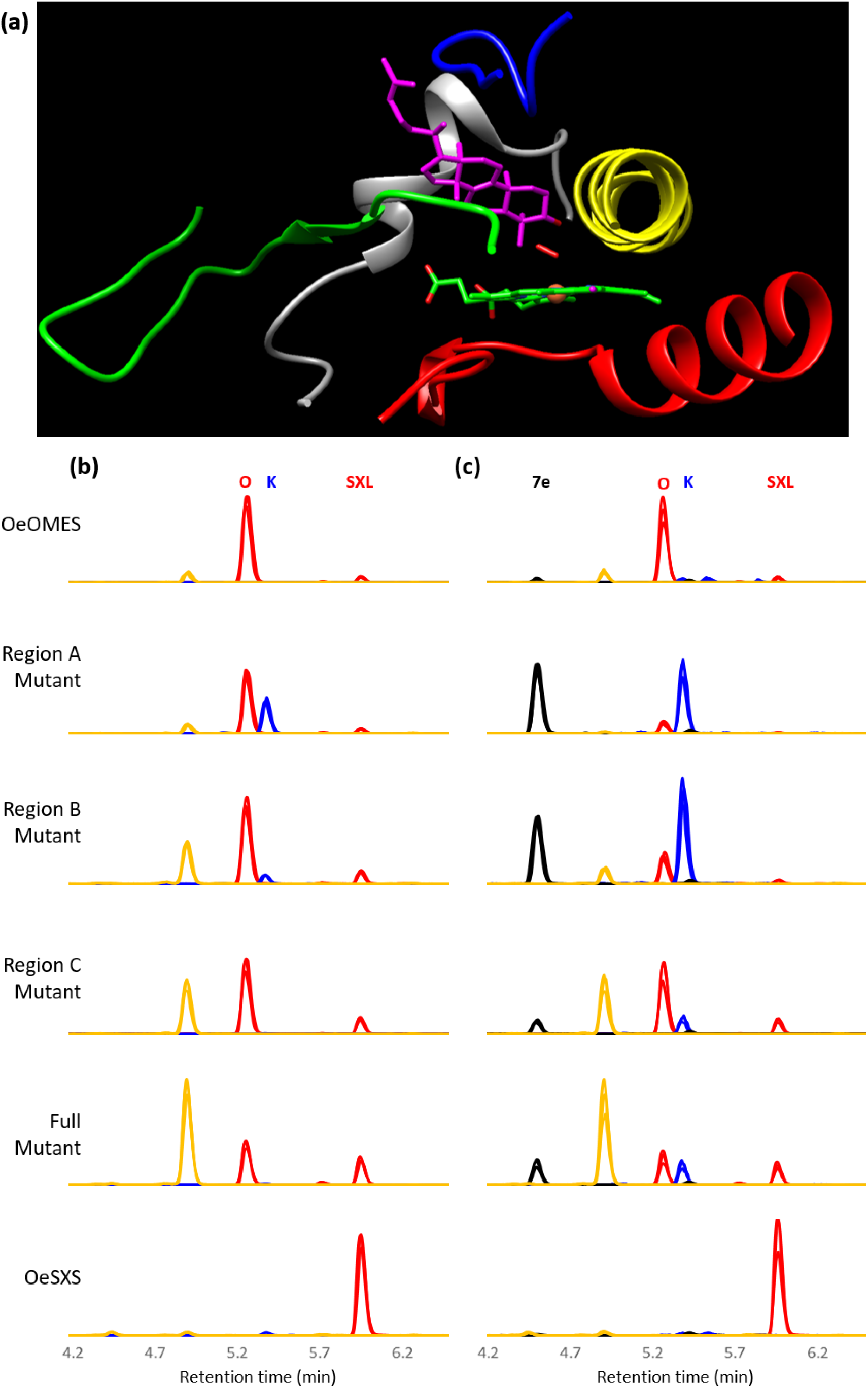
Extracted Ion Chromatograms (XIC) of biochemical assays of OeOMES mutants. (a) Homology model of OeOMES, using the backbone of Lanosterol 14-α demethylase (L14D; PDB 4LXJ; Monk et al., 2014) as a guide, showing the original ligands. Only regions of interest are shown, in colors: Gray (region A), Yellow (region B), Green (region C), Red (region D) and Blue (region E). XIC of the microsomal incubations with ketologanin (b) and 7-epi-loganin (c) are shown for selected mutants. Four channels are depicted, corresponding to the most abundant adduct of OME and secoxyloganin (red, [M-H]^−^ = 403.1240±0.05), ketologanin (blue, [M+FA-H]^−^ = 433.1346±0.05), 7-epiloganin (black, [M+FA-H]^−^ = 435.1503±0.05), and what we hypothesize to be the formic acid adduct of C_17_H_24_O_11_ (yellow, [C_17_H_24_O_11_+FA-H]^−^ = 449.1295±0.05). Intensities are scaled to the highest intensity of the corresponding channel in all incubations. Three replicates are shown, overlaid, for each mutant. O, oleoside; K, ketologanin; SXL, secoxyloganin.

Wild type OeOMES produces OME from 7-epi-loganin and ketologanin, with only trace amounts of secoxyloganin being formed. Production of secoxyloganin from ketologanin and 7-epiloganin increased in three mutants (Fig. 4): mutants in region B (4 amino acids switched), which is close to the Fe-ligand interaction; mutants in the region C (4 amino acids), a loop and sheet close to the substrate; and the fully switched mutant (22 amino acids). The latter had the highest production of secoxyloganin of the mutants consistently producing 75% as much secoxyloganin as OME from both 7-epiloganin and ketologanin, along with a decrease in OME production (Fig. 4). Notably, a new product was also observed in the secoxyloganin-producing mutants (Fig. 4, yellow chromatogram). This new product has a m/z corresponding to the formic acid adduct of an oxidation product of ketologanin, [C_17_H_24_O_11_+HCOO^−^]^−^.

Interestingly, the mutations in region A (5 amino acids), which is close to the substrate, opposite from the heme-coordinating region, had a strong negative effect in production of OME from 7-epiloganin (Fig. 4b), which is ameliorated when fed ketologanin (Fig. 4a). Similarly, the mutant in region E (3 amino acids), a loop physically close to region A and the substrate, when fed ketologanin produces as much OME as the wild type, but not when incubated with epi-loganin (Fig. S8). Mutations in regions D (6 amino acids), the heme-coordinating region, respectively, had no effect in any of the tested activities (Fig. S8, S9). Secologanin was also tested as a substrate to monitor overall oxidative activity. All mutants, regardless of the observed product profile with ketologanin, efficiently produced secoxyloganin from secologanin (Fig. S9).

### Analysis of expression data of olive iridoid pathway

To further explore the *in planta* properties of OeOMES and OeSXS, we analyzed the extensive publicly available expression data for olive. We separated the data in two sub groups: a study of 56 commercially cultivated varieties and wild olives (PRJNA525000, Gros-Balthazard *et al*., 2019); and five studies, comprising 47 datasets, targeting different biotic and abiotic stresses on olive (Leyva-Pérez *et al*., 2015; Jiménez-Ruiz *et al*., 2018; Jiménez-Ruiz *et al*., 2019). In the latter dataset, regardless of cultivar or stress, *OeISY* is always highly correlated with *OeOMES* (r = 0.67) and *OeSXS* (r = 0.58; Fig. S10). Interestingly, *OeISY* is negatively correlated (r = −0.77) with *OeSLS1*, providing further evidence that OeSLS1 does not have a role in iridoid related metabolism (Fig. S10). The studies showed that the only stressors that increase *OeOMES* expression are in roots (Picual cultivar) 24h after wounding (PRJNA256033), and in roots (Frantoio cultivar) two days after a challenge with *Verticillium dahliae* (Fig. S11). Detailed experiments have measured expression in inflorescence tissue of cultivated and wild olives (two botanical varieties of *O. europaea* subsp. *europaea*, var. *europaea* and *sylvestris*, respectively), to obtain insights into domestication of olive (study PRJNA525000, Gros-Balthazard *et al*., 2019). Fascinatingly, *ISY*, Iridoid Oxidase (*IO*) and *OMES* (12-fold increase in average) were highly expressed in inflorescences of commercial cultivars, while lacking expression in most wild olives (Fig. 5). In contrast, almost no expression of *OeSXS* was found in either commercial and wild olives, and no pattern can be discerned (Fig. 5d). In all datasets, *OeOMES* had consistently higher expression than *OeISY*.

**Fig. 5.**
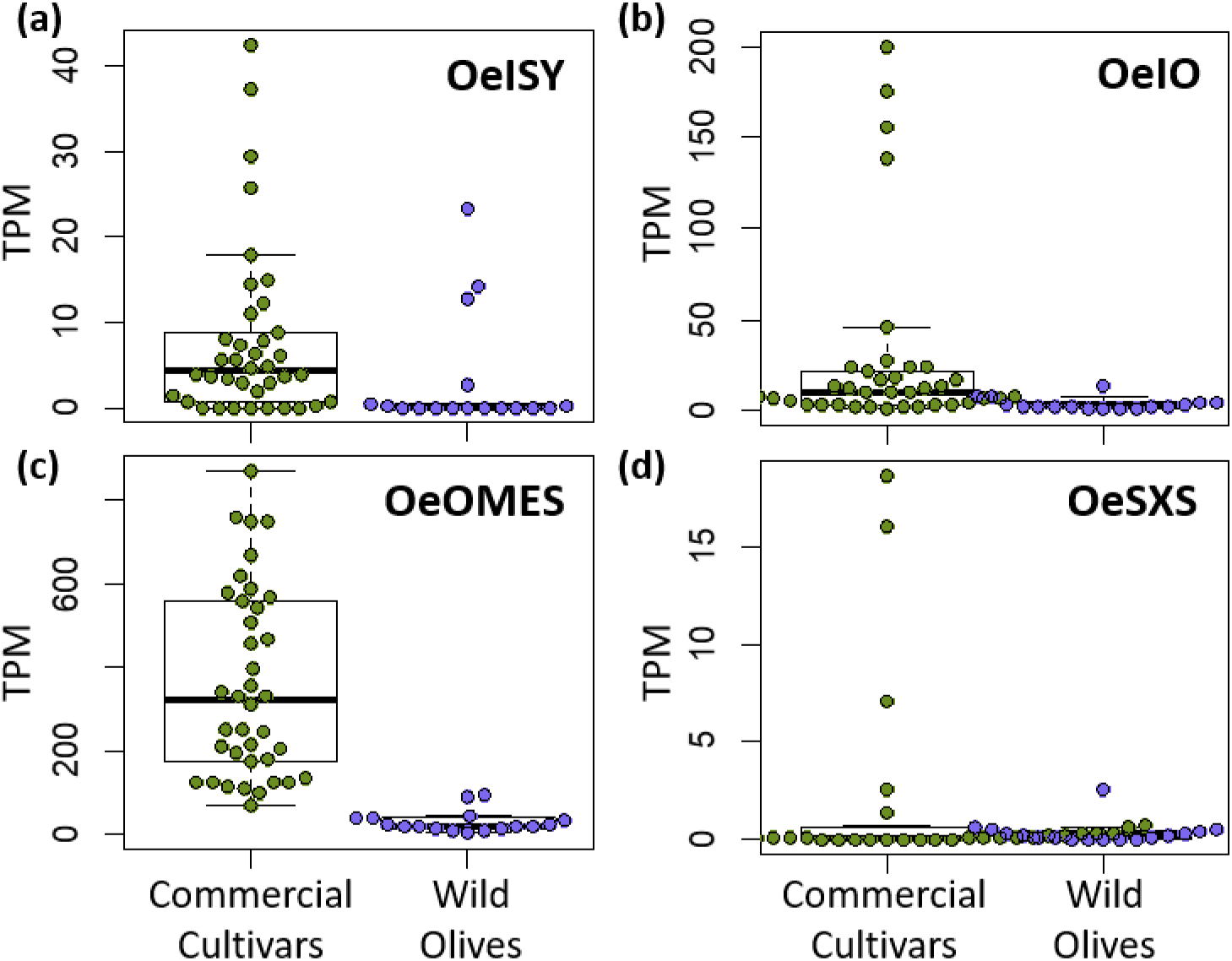
Expression levels of relevant transcripts. Swarmed boxplots are shown of the expression levels in inflorescences, in Transcript per Million (TPM), of 38 commercial cultivars (olive green) and 17 wild olives (light blue), of Iridoid Synthase (A), Iridoid Oxidase (B), Oleoside Methyl Ester Synthase (C) and Secoxyloganin Synthase (D). Raw data was obtained from (Gros-Balthazard *et al*., 2019) via NCBI SRA.

## Discussion

Here we report the discovery of the enzyme OeOMES, that catalyzes the oxidative cleavage leading to the major secoiridoid scaffold in olive, oleoside methyl ester (OME), the penultimate step of oleuropein biosynthesis. Previously, reported feeding studies in olive failed to identify the biosynthetic intermediates that lead to OME. These biochemical studies now show that OeOMES converts 7-epi-loganin via a two-step oxidation to form OME. We also identified a variant of this enzyme that catalyzes secoxyloganin biosynthesis, a metabolite that is also present in olive. Unexpectedly, secoxyloganin is also derived from 7-epi-loganin, and in *O. europaea* is not synthesized via secologanin, as we anticipated. This discovery highlights that secologanin synthase homologues have evolved to catalyze several oxidative cleavage reactions, as summarized in Fig. 6. Co-expression analysis shows that *OeISY*, *OeOMES* and *OeSXS* are co-regulated, while *OeSLS1* is negatively correlated to *OeISY*. Along with the fact that we detected no activity for OeSLS1, this supports the notion that OeSLS1 is not involved in iridoid biosynthesis. The kinetic experiments have shown that the oxidative cleavage of ketologanin by OeOMES is slightly slower, compared to ketologanin formation from 7-epiloganin by the same enzyme. Overall, however, the catalytic efficiencies for the two enzymes for both substrates are similar. Since only Oleaceae and select members of the Loasaceae (Weigend *et al*., 2000) produce oleoside-type secoiridoids, while also producing secologanin-type, we hypothesize that the emergence of the exocyclic olefin products is evolutionarily recent.

**Fig. 6.**
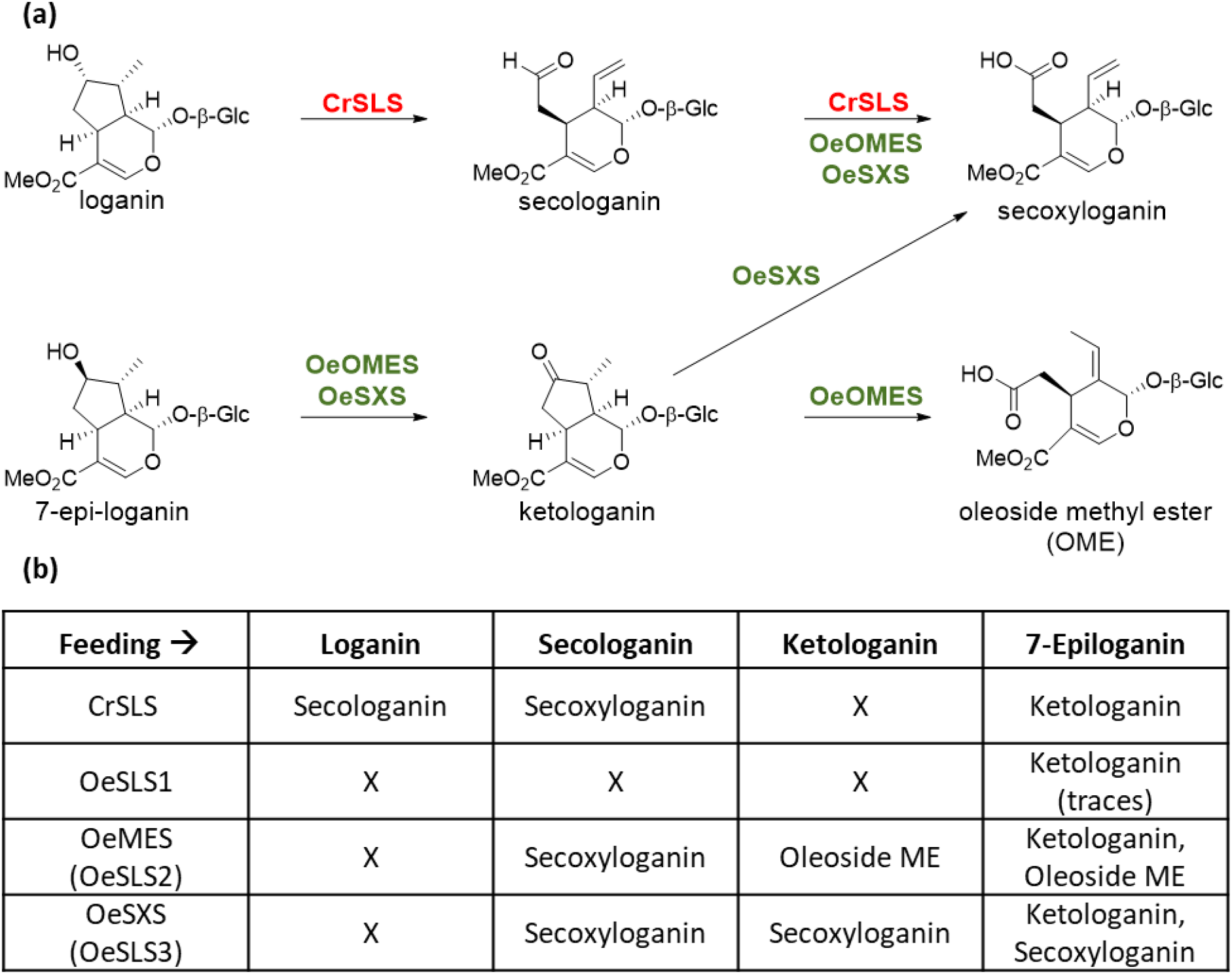
Summary of reactions producing biologically relevant metabolites. A graphical summary of reactions leading to biologically relevant metabolites is shown (a), along with a table summarizing the products of each enzyme, when incubated with different substrates.

These biochemical findings are largely consistent with previously published feeding experiments, particularly the thorough studies of Damtoft, Franzyk and Jensen (1993), in which deuterated iridoids were fed to *Syringa josikaea* and *Fraxinus excelsior*, both members of the Oleaceae. These prior studies demonstrated that while deoxyloganic acid was incorporated into secoiridoids in both *S. josikaea* (12.5%) and *F. excelsior* (11.5%), 8-epi-deoxyloganic acid was not metabolized by either, consistent with the substrate selectivity observed for OeOMES and OeSXS. Notably, Damtoft *et al*. reported incorporation of both loganic acid and 7-epi-loganic acid into the oleoside ligstroside (18% and 15%, respectively) (ligstroside is an analogue of oleuropein in which the oleoside methyl ester is derivatized with a tyrosol ester instead of a dihydroxyl phenyl ester). Although loganic acid was not turned over by OeOMES and OeSXS, it is possible that loganic acid could be oxidized to ketologanin by another plant oxidoreductase, and ketologanin could then be incorporated into ligstroside via OMES (Fig. 1).

CrSLS, OeOMES and OeSXS all oxidize secologanin to secoxyloganin, and all also oxidize 7-epiloganin to ketologanin. Notably, although the olive enzymes can turnover secologanin, we found no olive cytochrome P450 that generates secologanin, and there are no reports of secologanin in olive. Unexpectedly, OeOMES, which synthesizes the oleoside methyl ester, is able to oxidize secologanin. Given the lack of secologanin in olive tissues, it is likely that the ability of OeOMES and OeSXS to turnover secologanin to secoxyloganin is not a physiologically relevant activity. Similarly, neither 7-deoxyloganin, 7-epiloganin nor ketologanin have been detected in *C. roseus*, so the capability of CrSLS to turnover 7-epiloganin and 7-deoxyloganin is also likely not relevant *in planta*.

Although not all of these oxidation reactions may be physiologically relevant, they can shed light into the mechanism of these enzymes. We speculate that the stereochemistry of the hydroxyl group in position 7 of loganin determines the orientation of the substrate in the CrSLS binding site, such that the hydrogen of C10 can be abstracted by the iron cofactor, which ultimately leads to secologanin (Yamamoto *et al*., 2000; Fig. 7a). Formation of secoxyloganin by OeSXS could follow a similar mechanism (Fig. 7b), with the abstraction of the hydrogen of C10 ultimately leading to the ring opening. Conversely, when 7-deoxyloganin binds to CrSLS, we hypothesize that the substrate binds such that the hydrogen of carbon 7 reacts with the iron cofactor, leading to the formation of loganin (7*S*-OH) (Fig. S12b). Similarly, it appears that 7-epiloganin (7*R*-OH) also binds with the C7 H available to the iron cofactor, resulting in formation of ketologanin (Fig. S12c).

**Fig. 7.**
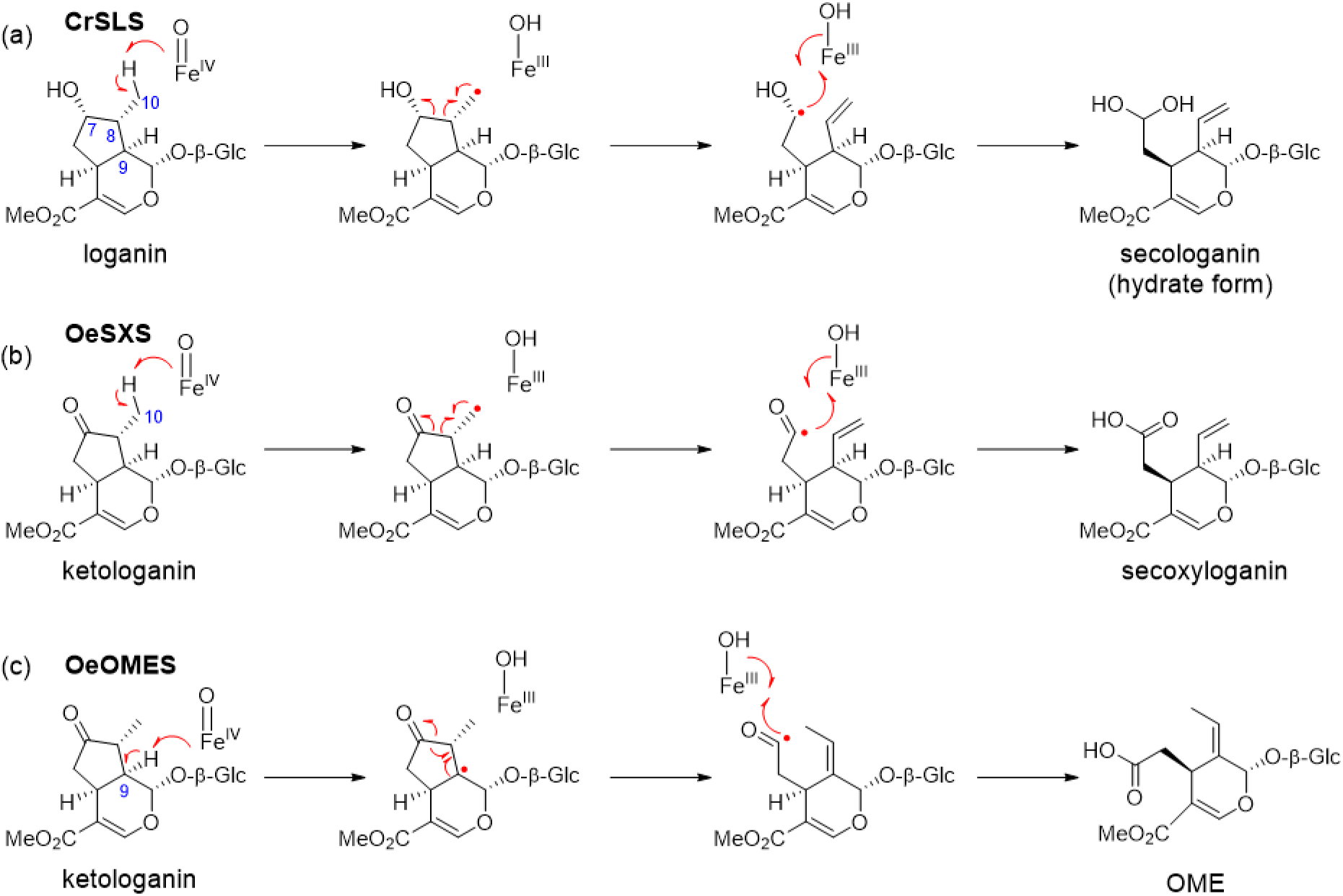
Mechanistic scenarios of secoiridoid formation. A summary of the proposed mechanistic scenarios of secologanin formation by CrSLS (a), secoxyloganin formation by OeSXS (b), and oleoside methyl ester (OME) formation by OeOMES (c).

It is likely that the oxidative cleavage of ketologanin by OeOMES follows a different mechanism than CrSLS. One mechanistic scenario would involve the abstraction of the hydrogen from Carbon 8 followed by rearrangement of the radical to Carbon 9, which could then open to form OME. Alternatively, the hydrogen of C9 could be abstracted directly (Fig. 7c). Notably, some of the mutants accumulated a side product, which although could not be isolated in sufficient quantity, displayed an m/z corresponding to a hydroxylated ketologanin derivative. We hypothesize that in these mutants (regions B and C, Fig. 4) one of these radical intermediates is quenched with a hydroxyl moiety to form this side product.

The mutational study of OeOMES provides some insights into which regions of the enzyme control substrate binding. Swaps of region A and region E led to a significant decrease in the oxidation, so these may be the major regions responsible for substrate coordination. Similarly, amino acid shifts in regions B and C caused a considerable increase in secoxyloganin production, revealing the role of these regions in product selectivity, possibly by modulating the position of the substrate relative to the activated iron species. The combination of all mutants, 22 amino acid shifts in total, led to a higher production of secoxyloganin, and a lower production of OME, likely a product of a synergy between the above-mentioned regions. Remarkably, inflorescences of commercial cultivars of olive show higher expression of genes in the oleuropein biosynthesis pathway, especially OeOMES, when compared to their wild counterparts. It has been hypothesized that domestication of olive was driven by gene expression, rather than genomic changes (Gros-Balthazard *et al*., 2019). It has also been reported that secoiridoid contents are mainly controlled by the cultivar genotype (Miho *et al*., 2018; Pérez *et al*., 2018; Deiana *et al*., 2019). Notably, published transcriptomic datasets for olive have shown that OeOMES expression is not substantially increased by environmental stresses, but is more highly expressed in domesticated versus wild olives.

For thousands of years, olive trees have been used to produce olive oil, and secoiridoids are a major effector in taste and quality. The expression profiles of OeOMES in the extensive transcriptomic data for olive points to a connection between domestication and iridoid metabolism. These newly identified genetic determinants of secoiridoid metabolism are of great value for breeding programs, and can help develop next-generation cultivars with ‘fine-tuned’ nutritional and taste characteristics.

## Acknowledgements

Our work was supported by the Alexander von Humboldt Foundation (Benke Hong), the Max-Planck-Gesellschaft and the Mint Consortium (Carlos Rodríguez and Sarah O’Connor). Further support derived from the European Union’s Horizon 2020 Research and Innovation Program Marie Sklodowska-Curie - Before Project (Grant Agreement No 645595) (Ornella Calderini, Luciana Baldoni, Valentina Passeri and Konstantinos Koudounas), the Italian National Research Council, Short-Term Mobility Program (Ornella Calderini) and the Rural Development Program of Umbria Region, 2014-2020 – Measure 16.2.1, Project INNO.V.O. (Ornella Calderini, Luciana Baldoni, Valentina Passeri), MIUR-DM n.856 del 10-10-2019 - Riparto Fondo Ordinario Enti di Ricerca - Anno 2019 - Assegnazione CNR per Progettualità di carattere Straordinario - Progetto “Economia Circolare (Green&CircularEconomy - GECE) (Ornella Calderini and Luciana Baldoni).

## Author Contributions

C.E.R.-L., O.C. and S.O.C designed experiments and wrote the manuscript; C.E.R.-L. and O.C. identified, cloned and characterized enzymes; C.E.R.-L. designed all mutants and performed all kinetic analyses; B.H. synthesized all standards and substrates; C.P and Y.N. determined the absolute configuration of the OME standard; K.K., V.P., L.B. and F.A. provided plant material and gene sequences and assisted with transcriptome analysis.

## Data Availability Statement

The data that support the findings of this study are publicly available or available from the corresponding author upon reasonable request.

