## Supplementary Files for "Two bi-functional cytochrome P450 CYP72 enzymes from olive (*Olea europaea*) catalyze the oxidative C-C bond cleavage in the biosynthesis of secoxy-iridoids - flavor and quality determinants in olive oil"

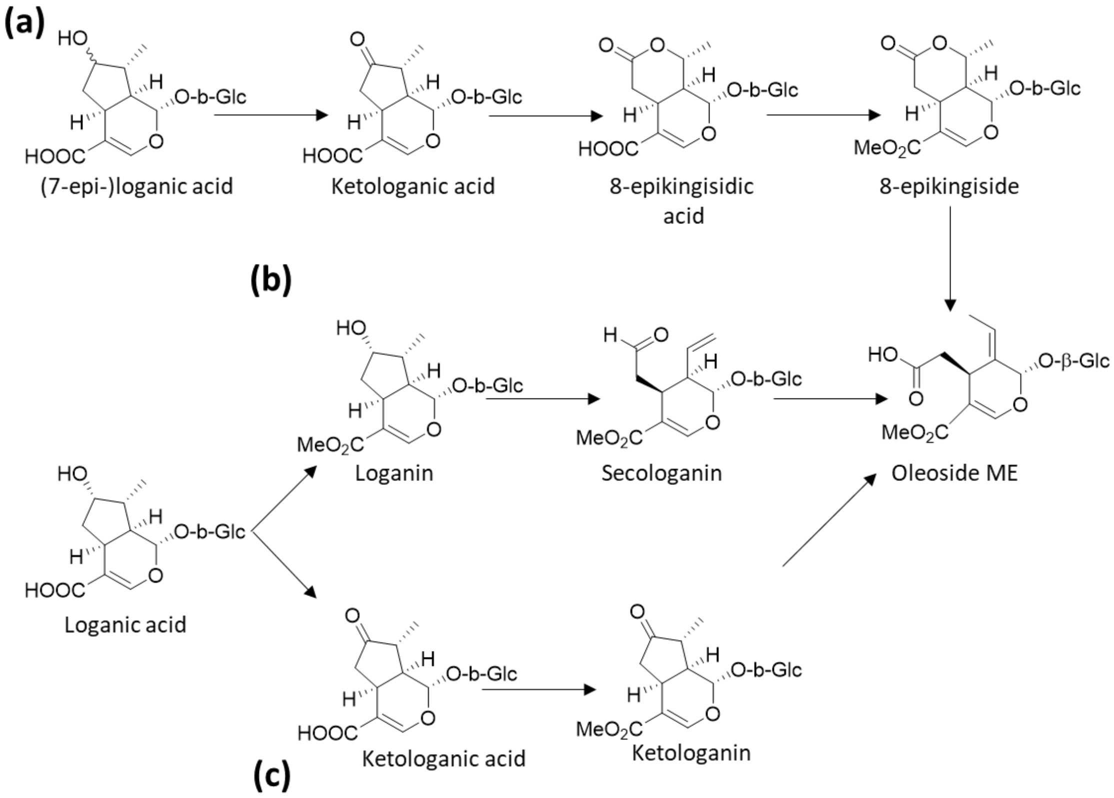


**Fig. S1 Previous hypotheses on the biosynthesis of secoiridoids in Olive**. Hypothesis differ on the immediate precursors: hypothesis (a), where 8-epikingiside is the immediate precursor (Inouye *et al.*, 1971,; Damtoft *et al.*, 1993); hypothesis (b), where it is secologanin (Inouye *et al.*, 1971,; Damtoft *et al.*, 1993; Alagna *et al.*, 2016); and hypothesis (c), being ketologanin (Alagna *et al.*, 2016). None of these hypotheses are supported by our results, but rather ketologanin is formed from 7-epiloganin, and then cleaved to form the oleoside methyl ester.

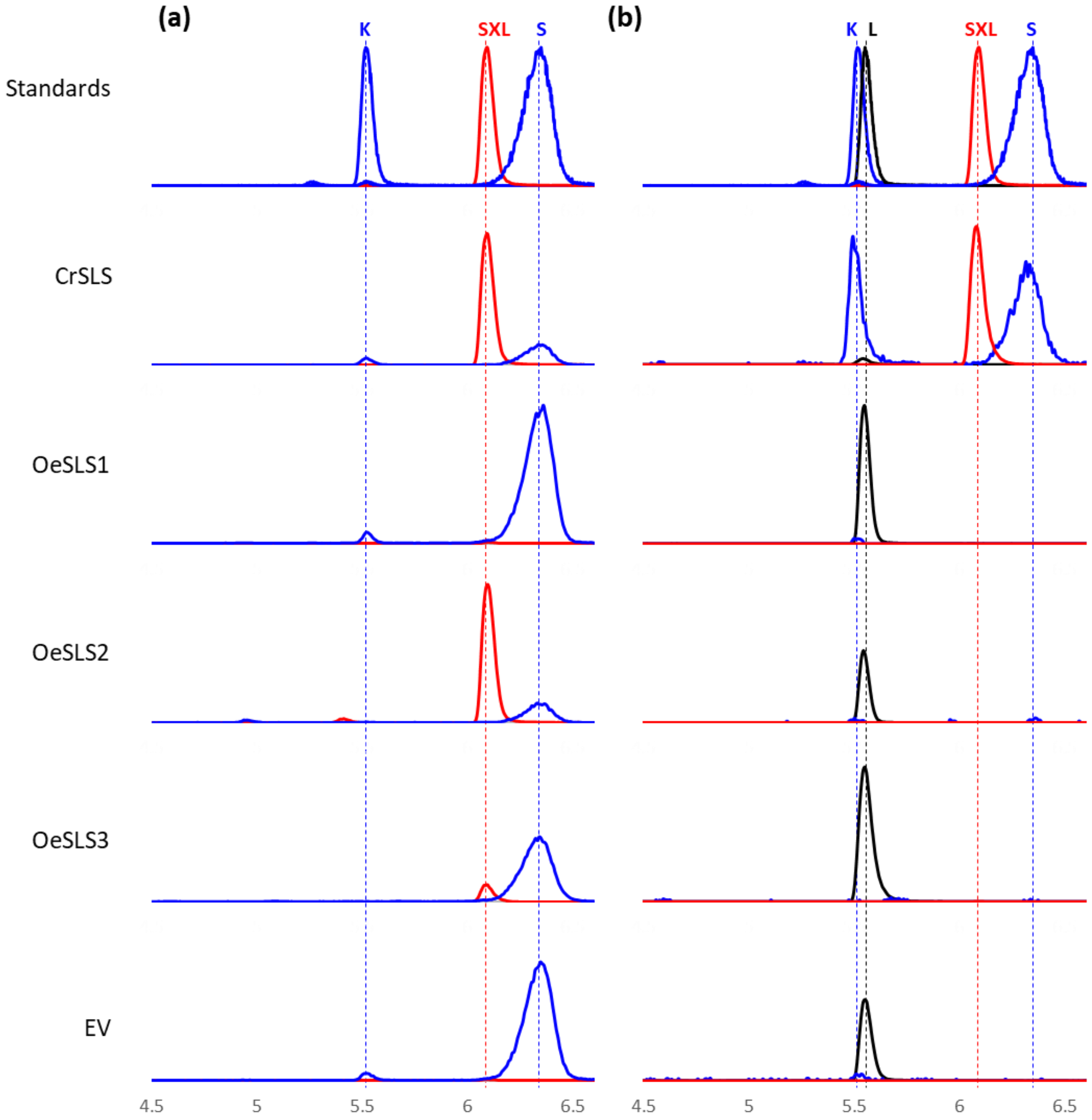


**Fig. S2 Extracted Ion Chromatograms (XIC) of microsome substrate assays.** XIC of the microsomal incubations with secologanin (a) and loganin (b) are shown. Three channels are depicted, corresponding to the most abundant adduct of secoxyloganin (red, $[M\text{-}H]^-$ = 403.1240±0.05), secologanin (blue, $[M\text{+}FA\text{-}H]^-$ = 433.1346±0.05) and loganin (black, $[M\text{+}FA\text{-}H]^-$ = 435.1503±0.05). Intensities are scaled to the highest intensity of the corresponding channel in all incubations. K, ketologanin; SXL, secoxyloganin; S, secologanin.

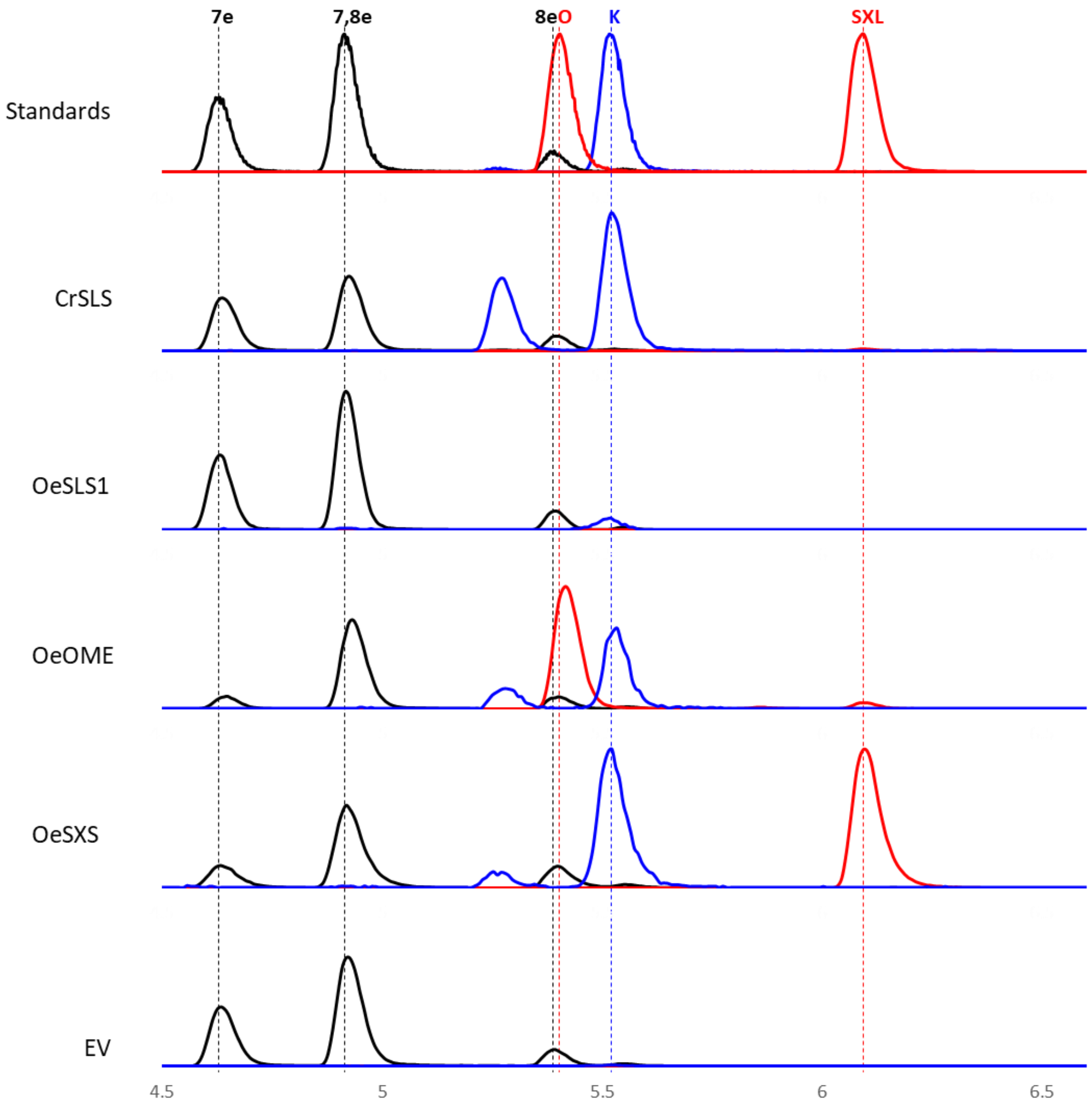


**Fig. S3 Extracted Ion Chromatograms (XIC) of microsome substrate assays.** XIC of the microsomal incubations with a mix of loganin epimers are shown. Three channels are depicted, corresponding to the most abundant adduct of OME and secoxyloganin (red, $[M-H]^-$ = 403.1240±0.05), ketologanin and secologanin (blue, $[M+FA-H]^-$ = 433.1346±0.05), and loganin epimers (black, $[M+FA-H]^-$ = 435.1503±0.05). Intensities are scaled to the highest intensity of the corresponding channel in all incubations. 7e, 7-epi-loganin; 7,8-e, 7,8-epi-loganin; 8e, 8-epi-loganin; O, oleoside; K, ketologanin; SXL, secoxyloganin.

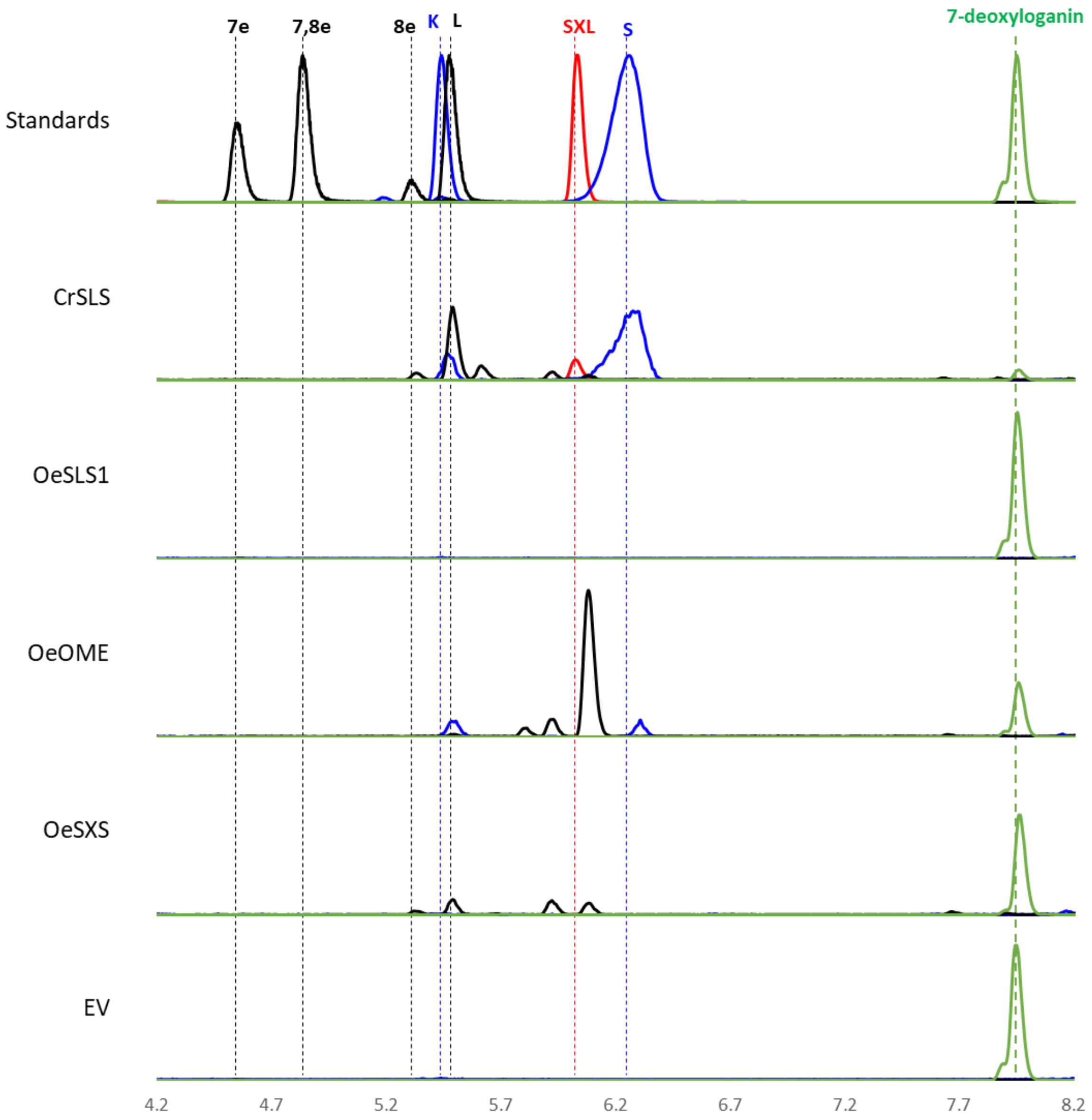


**Fig. S4 Extracted Ion Chromatograms (XIC) of microsome substrate assays.** XIC of the microsomal incubations with a mix of 7-deoxyloganin are shown. Four channels are depicted, corresponding to the most abundant adduct of secoxyloganin (red, $[M-H]^-$ = 403.1240±0.05), ketologanin and secologanin (blue, $[M+FA-H]^-$ = 433.1346±0.05), loganin epimers (black, $[M+FA-H]^-$ = 435.1503±0.05) and deoxyloganin (Green, $[M+FA-H]^-$ = 419.1553±0.05). Intensities are scaled to the highest intensity of the corresponding channel in all incubations. 7e, 7-epi-loganin; 7,8-e, 7,8-epi-loganin; 8e, 8-epi-loganin; L, loganin; K, ketologanin; SXL, secoxyloganin; S, secologanin.

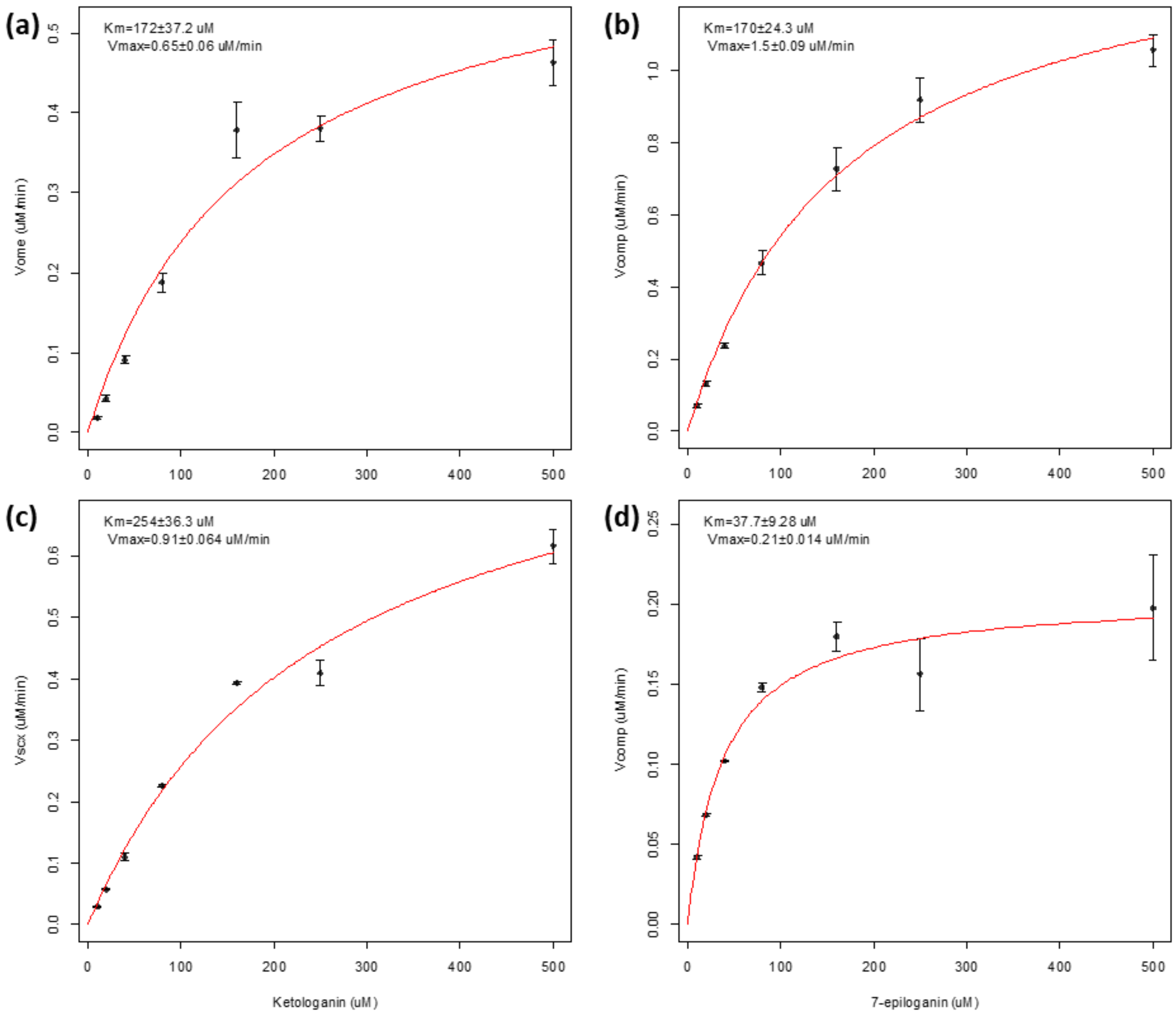


**Fig. S5 Kinetic modelling of enzyme activity.** Kinetics of production rate of OME by OeOMES (**a**) and secoxyloganin by OeSXS **(c)** with respect to ketologanin concentration; and scaled sum of velocities ($v_{comp}$) for OeOMES **(b)** and OeSXS **(d)** with respect to 7-epi-loganin concentration. Dots represent the average and error bars the standard error of three measurements, and the regression is shown in red.

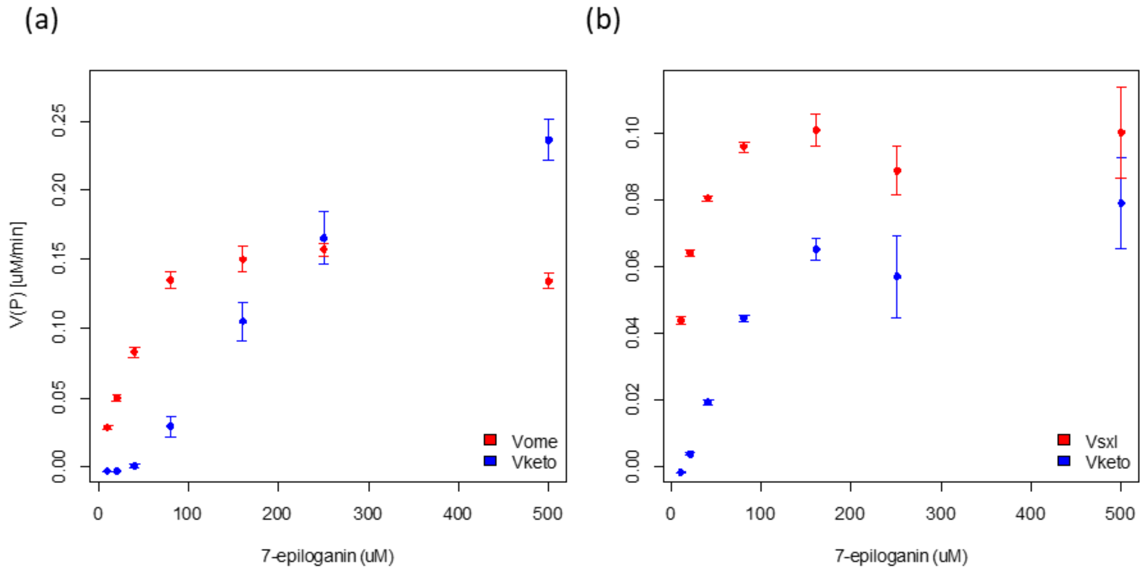


**Fig. S6 Individual enzyme activity.** Rates of production of OME (red) and ketologanin (blue) by OeOMES (**a**) and secoxyloganin (red) and ketologanin (blue) by OeSXS **(b)** with respect to 7-epi-loganin concentration. Dots represent the average and error bars the standard error of three measurements.

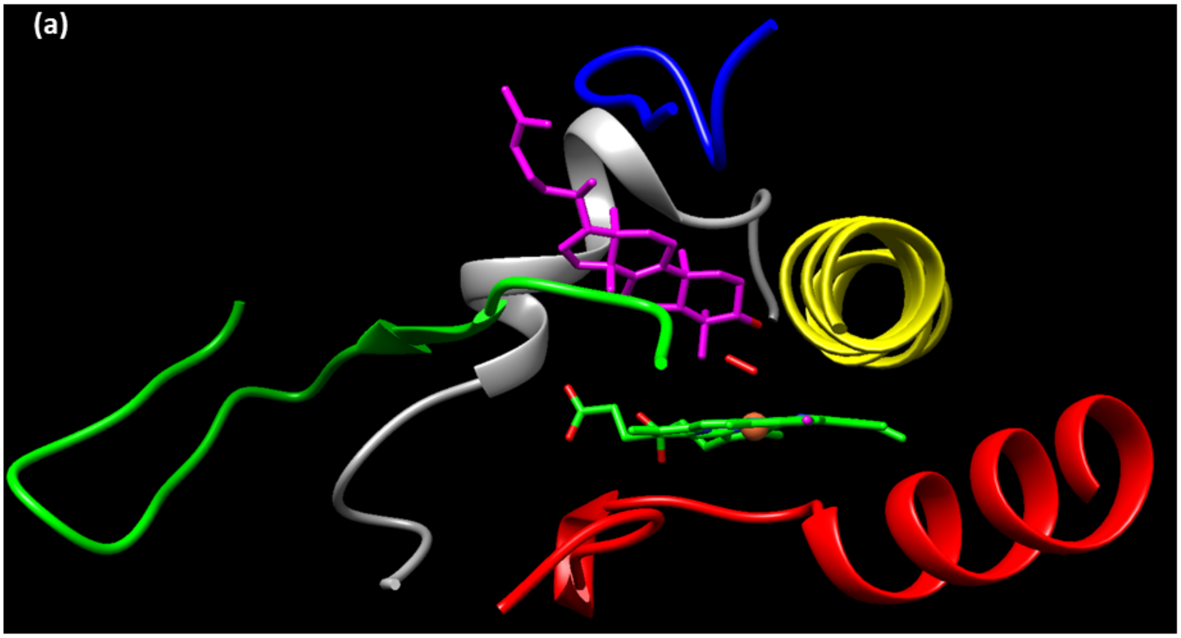


(b)

```
  1 MEILKTAISV VTIIIFTAAW RILKWLWLDP KKKERILRQQ GFKGNPYTFR
 51 RLLFGDEKEI QKIHAEAWSK PVNIRDEVQT GKRAAPFIFR TYEKYGKKAF
101 VWAGLRPKVF IMEPDHMKTI FFNHSTFQKN FKVTNSVVQE LITGIIRFEG
151 EEWSKRRTIM SPKFQLEKLK QMIPLVLKCS DQVIIEWKKL VSNSKDGSYT
201 LDVCHDIEEL ISGVTSQFLF GIDYAKDKEI FHLITQLSEL TKQATKVSNL
251 PGSKFLPTET NRKSKRLTKE LHQRLYKLMC ERKKAIKEGK VVEDNVFNML
301 LESEIANNQD EMIGHMKGFV FNSHDTTAFV LVWNLILLCI YSEWQDRARE
351 EVFRVFGNRR PDYEGLSQLK VLPMFMNEVL RLYPPLVELS RFLEEEIKLG
401 EYTLPADIQV IMPTILVHRD PEFWGEDANE FKPERFAEGV LKATNGQAVF
451 FPFAWGPRIC IGYNMALLQV KLVLADLLRN FSFEISPTYE HAPRVVFTQQ
501 PQYGAPIILR NLN
```

**Fig. S7 Regions of Interest**. (a) Homology model of OeOMES, using the backbone of Lanosterol 14-α demethylase (L14D; PDB 4LXJ; Monk et al., 2014) as a guide. Only the regions of interest, closest to the original substrate in the model of L14D, are shown. (b) OeOMES aminoacid sequence, with the regions of interest highlighted with the corresponding colors: Gray (region A), Yellow (region B), Green (region C), Red (region D) and Blue (region E).

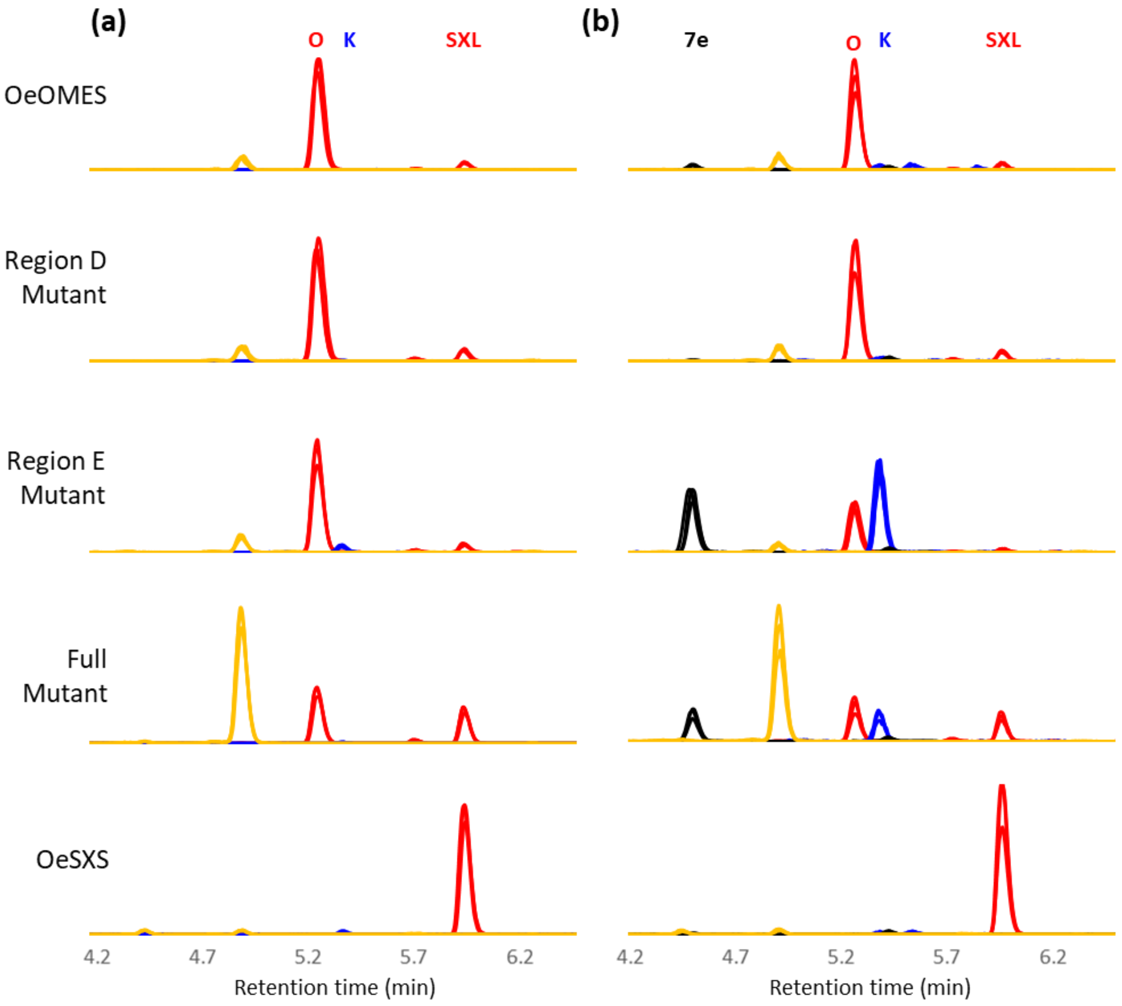


**Fig. S8 Extracted Ion Chromatograms (XIC) of microsome substrate assays.** XIC of the microsomal incubations with ketologanin (a) and 7-epi-loganin (b) are shown for selected mutants. Four channels are depicted, corresponding to the most abundant adduct of OME and secoxyloganin (red, $[M\text{-}H]^-$ = 403.1240±0.05), ketologanin (blue, $[M\text{+}FA\text{-}H]^-$ = 433.1346±0.05), 7-epiloganin (black, $[M\text{+}FA\text{-}H]^-$ = 435.1503±0.05), and oxidized ketologanin (yellow, $[C_{17}H_{24}O_{11}\text{+}FA\text{-}H]^-$ = 449.1295±0.05). Intensities are scaled to the highest intensity of the corresponding channel in all incubations. Three replicates are shown, overlaid, for each mutant. O, oleoside; K, ketologanin; SXL, secoxyloganin.

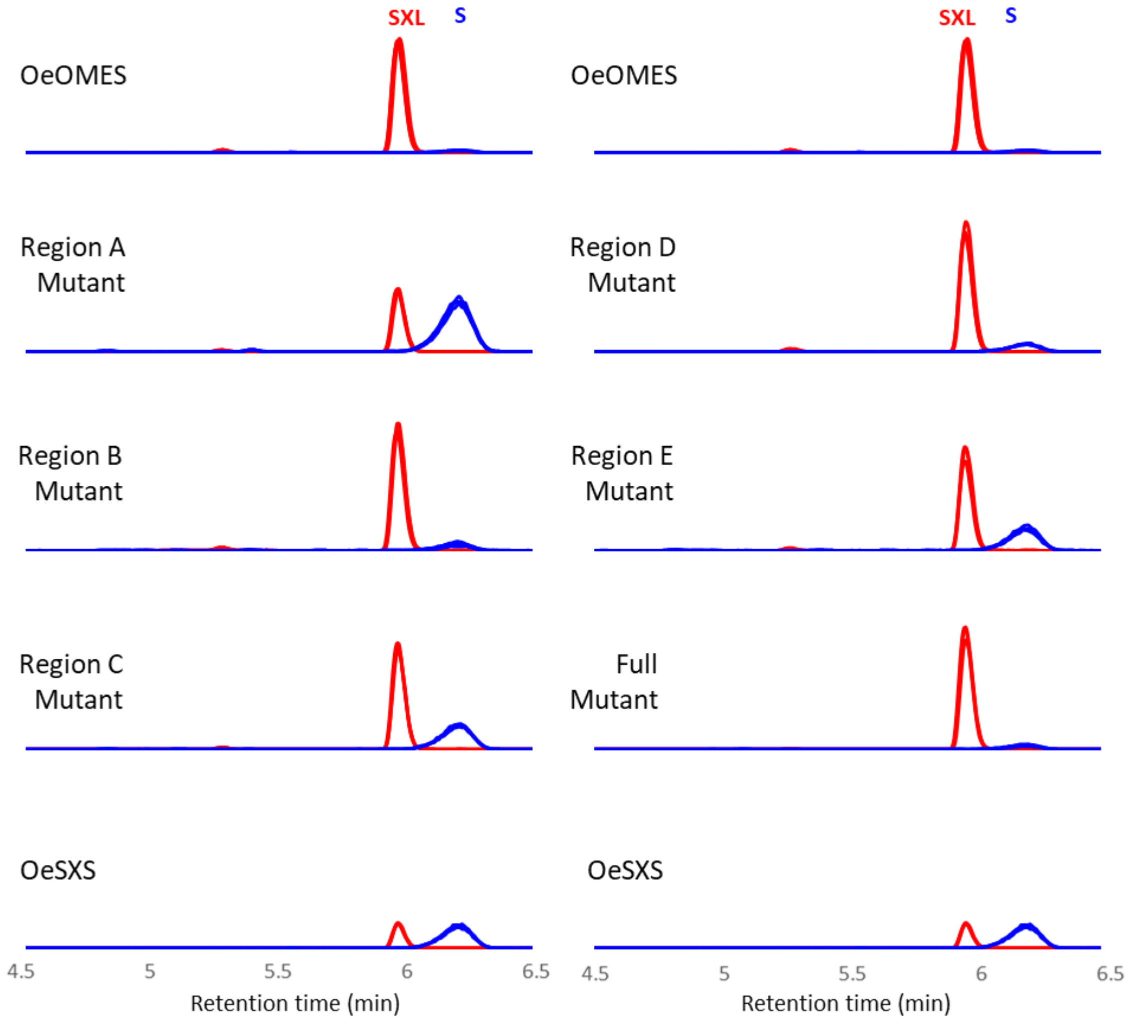


**Fig. S9 Extracted Ion Chromatograms (XIC) of microsome substrate assays.** XIC of the microsomal incubations with secologanin are shown for all mutants. Two channels are depicted, corresponding to the most abundant adduct of secoxyloganin (red, [M-H]- = 403.1240±0.05), and ketologanin (blue, [M+FA-H]- = 433.1346±0.05). Intensities are scaled to the highest intensity of the corresponding channel in all incubations. Three replicates are shown, overlaid, for each mutant. SXL, secoxyloganin; S, secologanin.

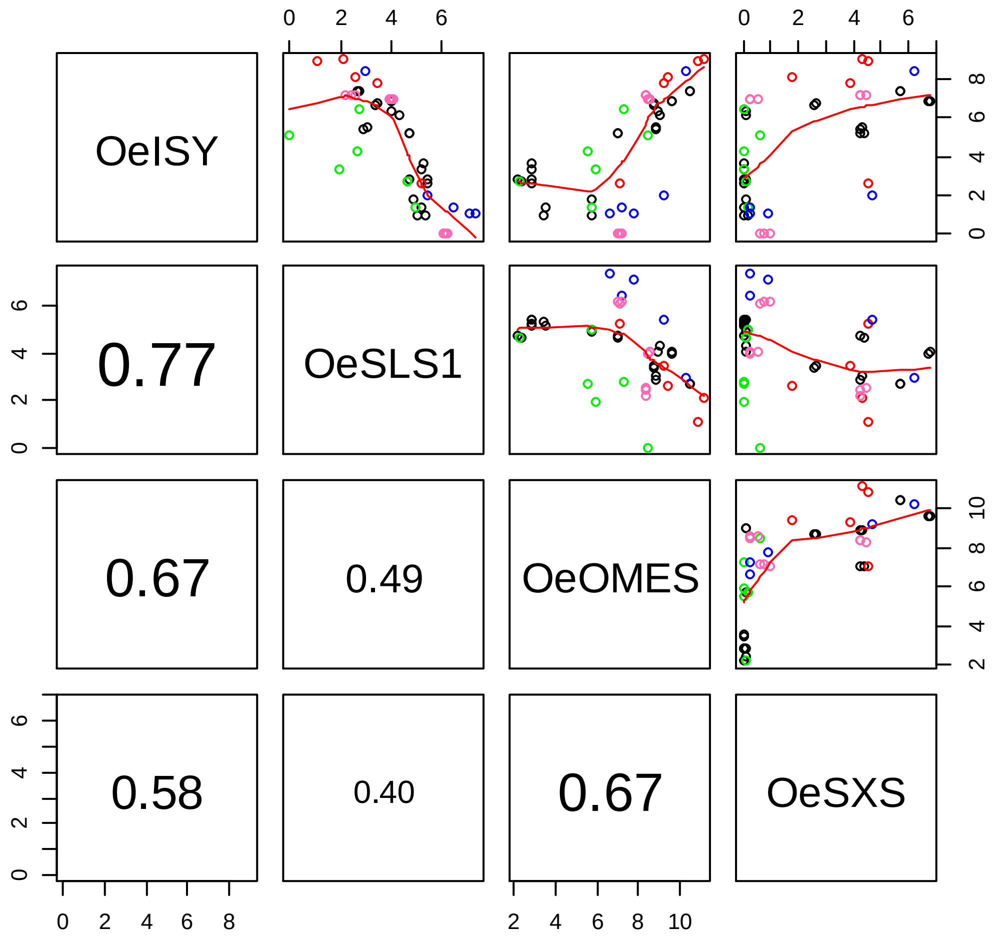


**Fig. S10 Coexpression Analysis**. A correlogram is shown of the coexpression of *OeISY*, *OeSLS1*, *OeOMES* and *OeSXS* in different BioProjects. The upper diagonal shows scatter plots of the logarithm, base 2, of the expression in Transcripts Per Million (TPM) of each gene, with a lowess regression on the data (red line). The lower diagonal shows the absolute of Pearson's product-moment correlation coefficient. Each point corresponds to an RNA-seq run of projects PRJNA256033 (black), PRJNA378602 (green), PRJNA401310 (red), PRJNA514943 (blue), and PRJNA596876 (pink)

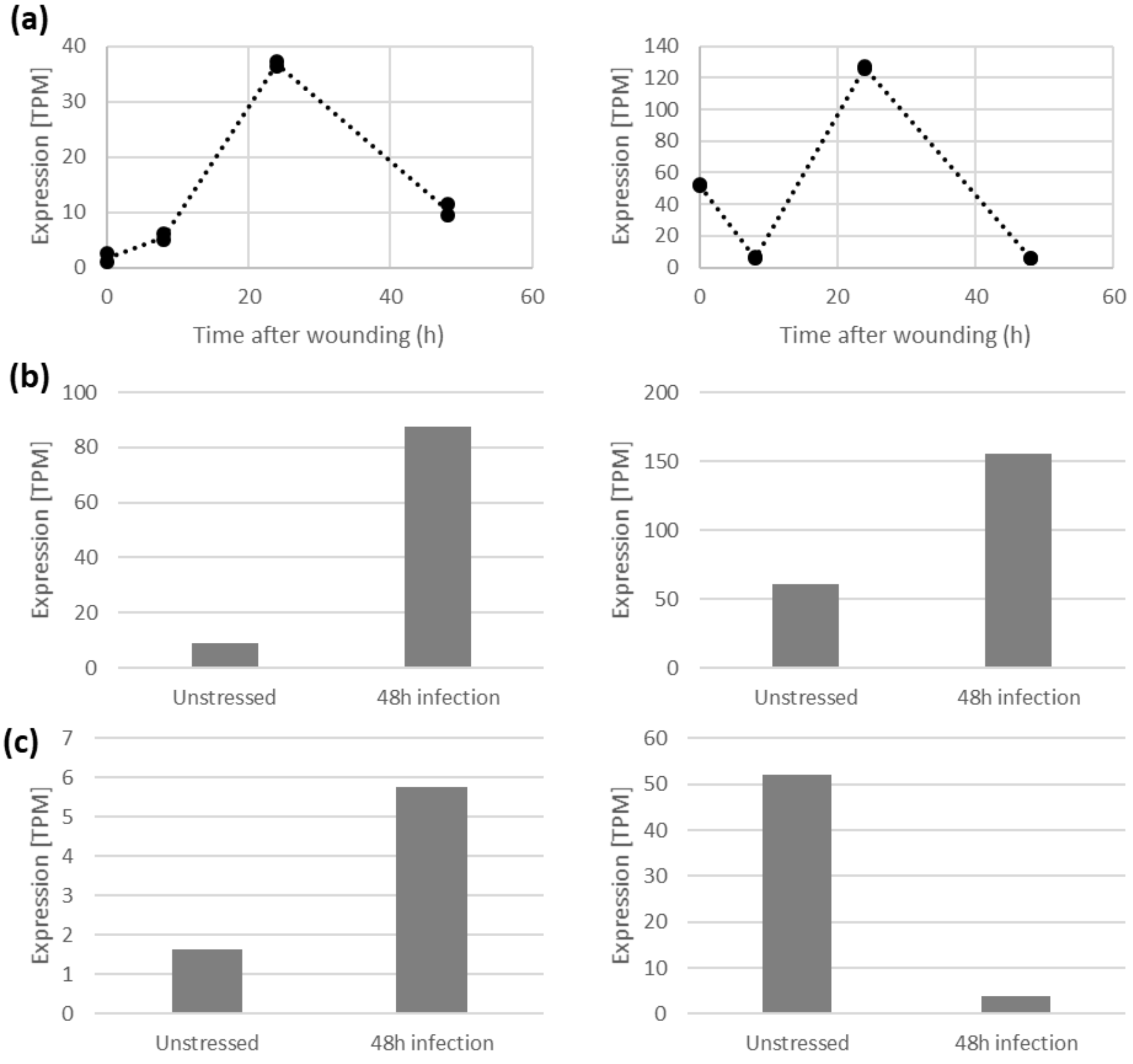


**Fig. S11 Effect of different stresses in OeISY and OeOMES expression**. Expression is shown for OeISY (left) and OeOMES (right) Picual roots after mechanical wounding (a), and Frantoio (b) and Picual (c) roots after infection with *Verticillium dahliae*. The last two experiments, (b) and (c) had a single replicate, while (a) was done in duplicate. For the latter, each point is a replicate.

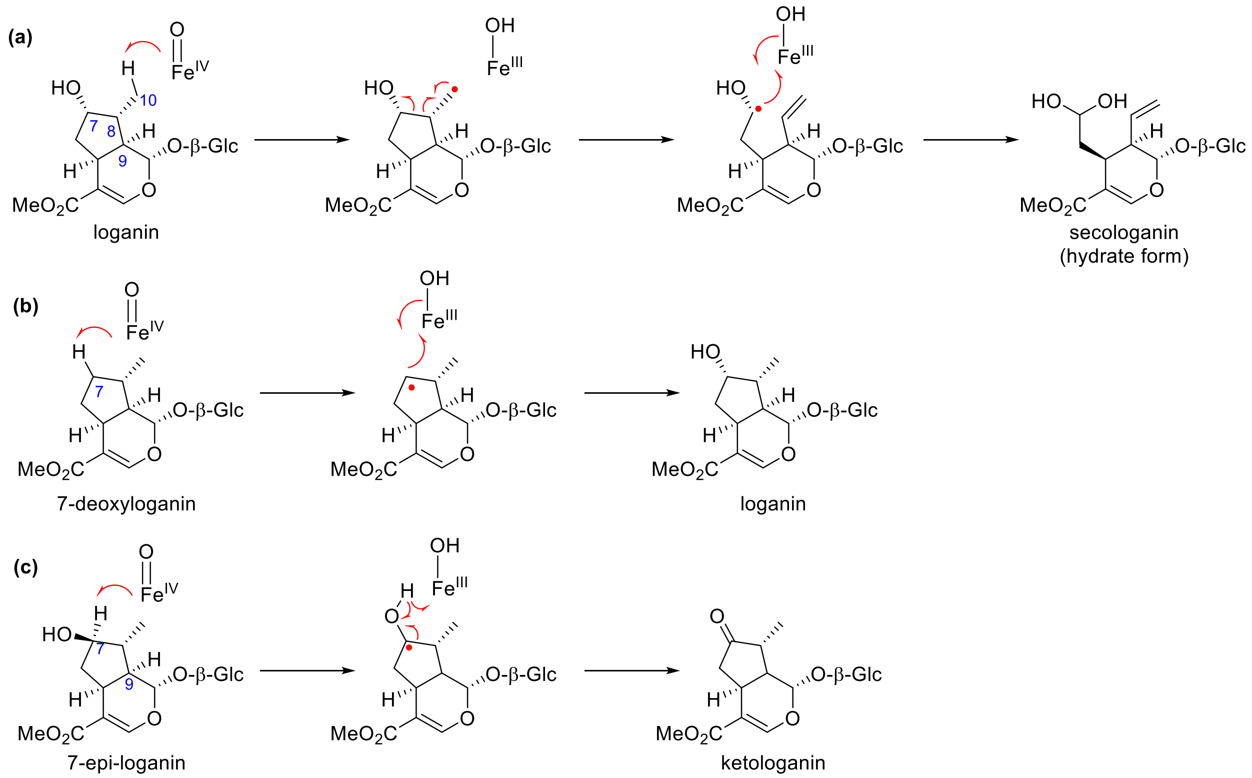


**Fig. S12 Proposed mechanisms of CrSLS catalyzed reactions**. A summary of the proposed mechanisms for production of secologanin (a), loganin (b) and ketologanin (c) by CrSLS. Mechanism (a) was adapted from (Yamamoto *et al.*, 2000).

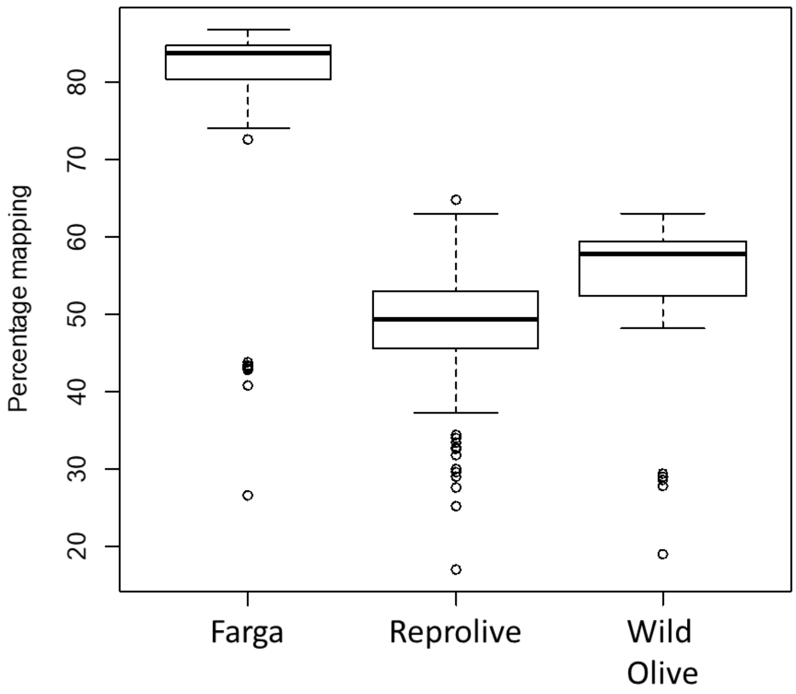


**Fig. S13 Mapping against available transcriptomes**. Percentage mapping of the 112 runs analyzed in this article against different available transcriptomes.

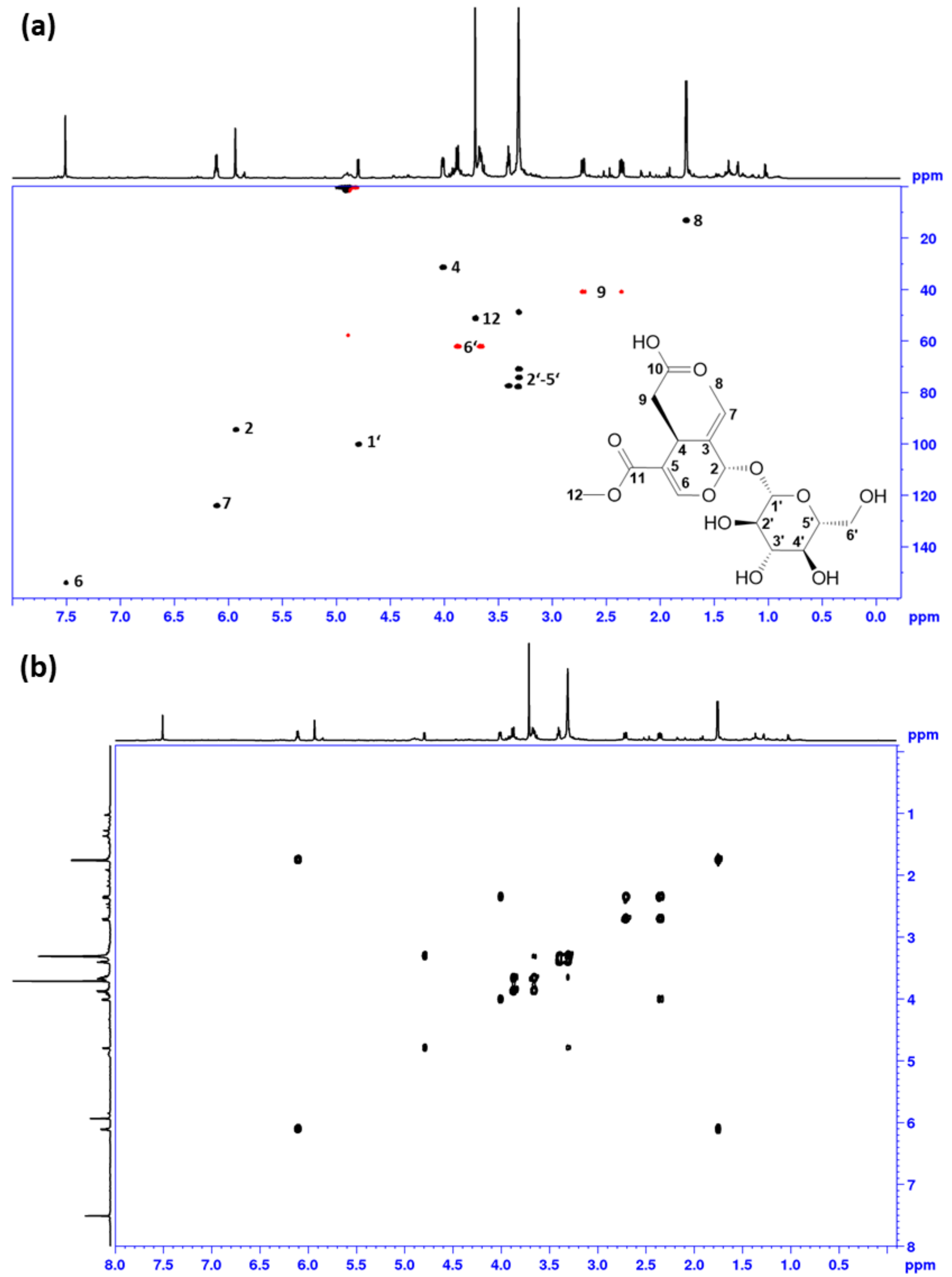


**Fig. S14 NMR spectra of the Oleoside Methyl Ester standard**. The $^{1}$H-$^{13}$C HSQC (a) and $^{1}$H-$^{1}$H DQFCOSY (b) of oleoside methyl ester are shown.

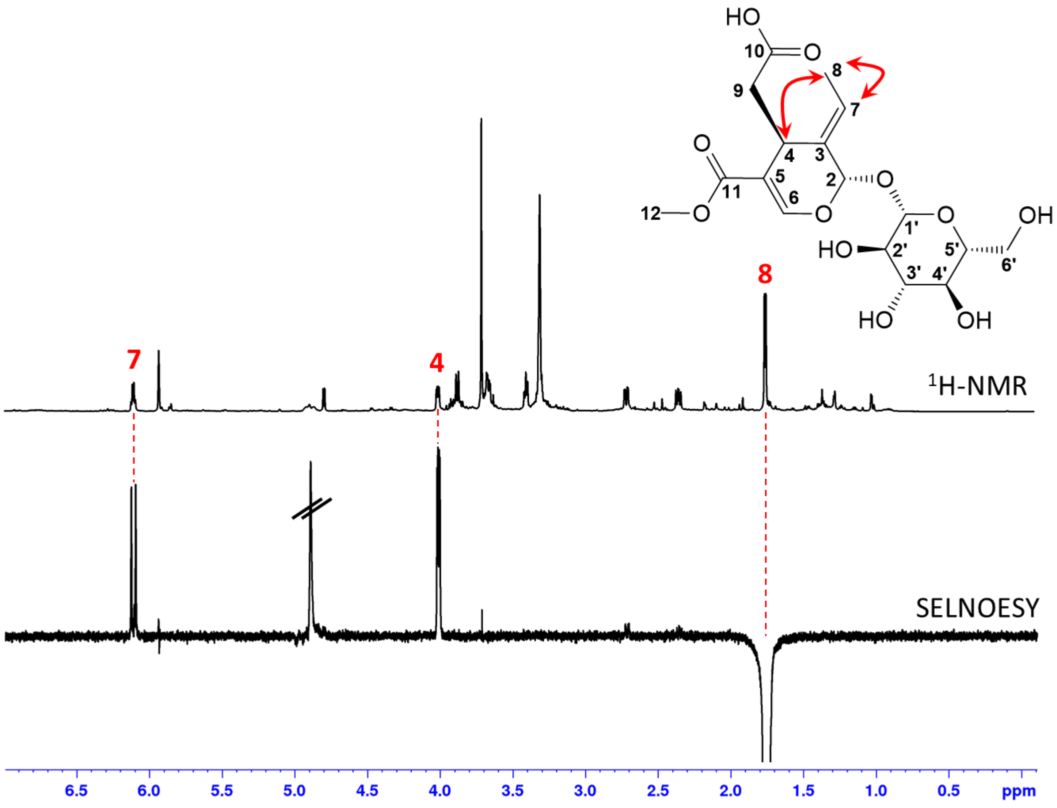


**Fig. S15 Confirmation of the Oleoside Methyl Ester configuration**. The $^1$H-NMR and selective NOESY spectra (mixing time of 600 ms) of oleoside methyl ester are shown. As transmitter frequency 1.785 ppm/1250.5 Hz was selected (methyl group in position 8). These spectra confirm the exocyclic olefinic bond is in the E configuration.

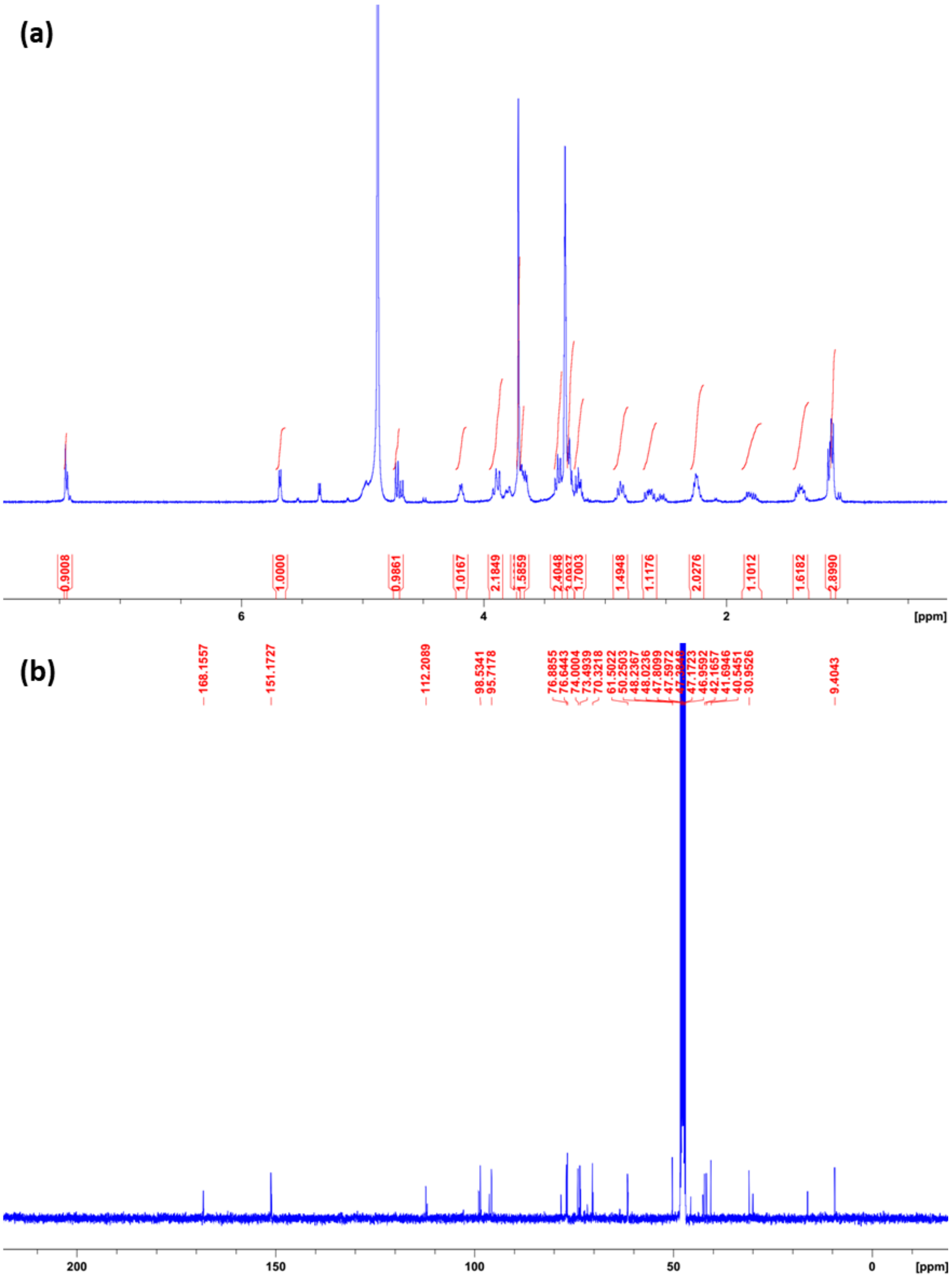


**Fig. S16 NMR spectra of 7-epi-8-epi-loganin**. The [1]H (a) and [13]C-NMR (b) spectrum of 7-*epi*-8-*epi*-loganin is shown.

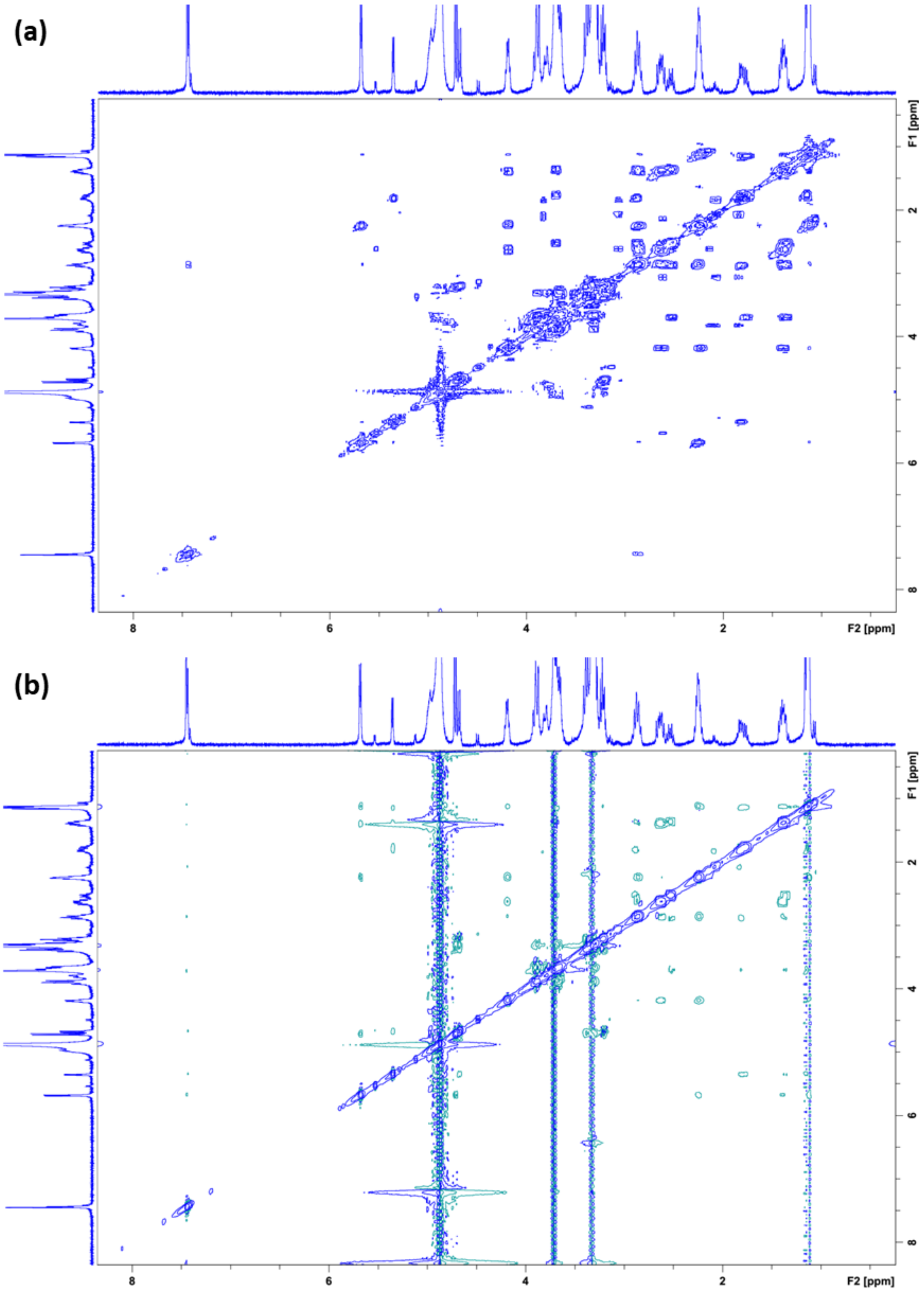


**Fig. S17 COSY and NOESY spectra of 7-epi-8-epi-loganin**. The $^{1}H$-$^{1}H$ COSY (a) and NOESY (b) spectrum of 7-*epi*-8-*epi*-loganin is shown.

**Table S1 Amino acid shifts in each of the selected regions.**

| Mutant | Amino acid shifts |
|---|---|
| Region A | 123 N→K<br>126 T→N<br>136 S→N<br>137 V→I<br>140 E→D |
| Region B | 320 V→F<br>323 S→A<br>329 F→V<br>330 V→L |
| Region C | 386 L→V<br>392 F→L<br>393 L→V<br>404 L→I |
| Region D | 450 F→Y<br>454 A→G<br>459 I→V<br>463 Y→Q<br>465 M→F<br>470 V→A |
| Region E | 494 R→H<br>496 V→I<br>499 Q→L |

**Notes S1 Steady-State kinetics of a coupled reaction.**

Assuming that enzyme regeneration by the CPR is much quicker than product oxidation, which is reasonable under excess NADPH, we can assume we have a coupled reaction of the type:

$$S + E \underset{k_{-1}}{\overset{k_1}{\leftrightarrow}} C_1 \overset{k_2}{\rightarrow} C_2 \overset{k_4}{\rightarrow} E + P \qquad \text{(Eq. S1.1)}$$

Where S is 7-epiloganin, E is the enzyme, $C_1$ is the enzymatic complex of Enzyme and 7-epiloganin, $C_2$ is the enzymatic complex of enzyme and ketologanin, and P is the final product (either secoxyloganin or the Oleoside Methyl Ester). Applying the law of mass action:

$$\frac{dC_1}{dt} = v_{C1} = k_1 SE - k_{-1}C_1 - k_2 C_1 \qquad \text{(Eq. S1.2)}$$

$$\frac{dC_2}{dt} = v_{C2} = k_2 C_1 - k_4 C_2 \qquad \text{(Eq. S1.3)}$$

$$\frac{dP}{dt} = v_P = k_4 C_2 \qquad \text{(Eq. S1.4)}$$

Assuming quasi-steady-state, $v_{C1} = 0$, and thus, substituting in Eq. S1.2:

$$v_{C1} = k_1 SE - (k_{-1} + k_2)C_1 = 0$$

$$\frac{(k_{-1} + k_2)C_1}{k_1} = K_{MS}C_1 = SE \qquad \text{(Eq. S1.5)}$$

Where $K_{MS} = \frac{(k_{-1}+k_2)}{k_1}$; the Michaelis-Menten constant of epiloganin. By mass balance, the free enzyme equals the initial concentration, minus the complexed enzyme: $E = E_0 - C_1 - C_2$; then:

$$K_{MS}C_1 = SE_0 - SC_1 - SC_2$$

$$(K_{MS} + S)C_1 = SE_0 - SC_2$$

$$C_1 = \frac{SE_0}{(K_{MS} + S)} - \frac{SC_2}{(K_{MS} + S)} \qquad \text{(Eq. S1.6)}$$

Now, substituting in (Eq. S1.3):

$$v_{C2} = k_2 C_1 - k_4 C_2 = \frac{k_2 S E_0}{(K_{MS} + S)} - \frac{k_2 S C_2}{(K_{MS} + S)} - k_4 C_2$$

And substituting with (Eq. S1.4):

$$v_{C2} = \frac{k_2 S E_0}{(K_{MS} + S)} - \frac{k_2 S C_2}{(K_{MS} + S)} - v_P$$

Since we can measure$v_P$, the secoxyloganin or OME production rate, and $v_{C2}$, given than the enzyme and ketologanin dissociate during the methanol extraction, we can group:

$$v_{C2} + v_P = \frac{k_2 S E_0}{(K_{MS} + S)} - \frac{k_2 S C_2}{(K_{MS} + S)}$$

$$v_{C2} + v_P = \frac{k_2 S}{(K_{MS} + S)}(E_0 - C_2) = \frac{k_2 E_0 S}{(K_{MS} + S)}\left(1 - \frac{C_2}{E_0}\right)$$

Given the maximum velocity of $C_2$ formation is $V_S = k_2 E_0$, and by (Eq. S1.4) $C_2 = \frac{v_P}{k_4}$ then:

$$v_{C2} + v_P = \frac{V_S S}{(K_{MS} + S)}\left(1 - \frac{v_P}{k_4 E_0}\right)$$

$$\frac{v_{C2} + v_P}{\left(1 - \frac{v_P}{k_4 E_0}\right)} = \frac{V_S S}{(K_{MS} + S)}$$

Since we can measure the maximum velocity of secoxyloganin or OME formation by assaying microsomal preparations with ketologanin, then we can obtain $k_4 E_0$ as the $V_X = k_4 E_0$, as long as the initial amount of enzyme is the same in both experiments. Then:

$$\frac{v_{C2} + v_P}{\left(1 - \frac{v_P}{V_X}\right)} = \frac{V_S S}{(K_{MS} + S)} \qquad \text{(Eq. S1.7)}$$

Which is equivalent to (Eq. 1) in the main text, when $v_x = 0$, since in this model there is no dissociation of ketologanin from the enzyme complex.

**Notes S2 Steady-State kinetics of a dissociative, sequential reaction.**

Assuming that enzyme regeneration by the CPR is much quicker than product oxidation, which is reasonable under excess NADPH, we can assume we have a sequential reaction of the type:

$$S + E \underset{k_{-1}}{\overset{k_1}{\leftrightarrow}} C_1 \overset{k_2}{\rightarrow} X + E \quad \text{(Eq. S2.1)}$$

$$X + E \underset{k_{-3}}{\overset{k_3}{\leftrightarrow}} C_2 \overset{k_4}{\rightarrow} P + E \quad \text{(Eq. S2.2)}$$

Where S is 7-epiloganin, E is the enzyme, $C_1$ is the enzymatic complex of Enzyme and 7-epiloganin, X is ketologanin, $C_2$ is the enzymatic complex of enzyme and ketologanin, and P is the final product (either secoxyloganin or the Oleoside Methyl Ester). By mass balance:

$$S_0 = S + C_1 + X + C_2 + P$$

Then:

$$\frac{dS_0}{dt} = \frac{dS}{dt} + \frac{dC_1}{dt} + \frac{dX}{dt} + \frac{dC_2}{dt} + \frac{dP}{dt} = 0$$

$$v_S + v_{C1} + (v_X + v_{C2}) + v_P = 0 \quad \text{(Eq. S2.3)}$$

Since we can measure $v_P$, and the sum of $(v_X + v_{C2})$, given than the enzyme and ketologanin dissociate during the methanol extraction, we can group (Eq. S2.3) as:

$$(v_X + v_{C2}) + v_P = -(v_S + v_{C1}) \quad \text{(Eq. S2.4)}$$

Applying the law of mass action:

$$v_S = -k_1 SE + k_{-1} C_1 \quad \text{(Eq. S2.5)}$$

$$v_{C1} = k_1 SE - k_{-1} C_1 - k_2 C_1 \quad \text{(Eq. S2.6)}$$

$$v_P = k_4 C_2 \quad \text{(Eq. S2.7)}$$

Then, adding (Eq. S2.5) + (Eq. S2.6):

$$(v_S + v_{C1}) = -k_2 C_1 \qquad \text{(Eq. S2.8)}$$

And substituting in (Eq. S2.4)

$$(v_X + v_{C2}) + v_P = -(v_S + v_{C1}) = k_2 C_1 \qquad \text{(Eq. S2.9)}$$

Assuming quasi-steady-state in the beginning of the incubation, $v_{C1} = 0$, and thus, substituting in Eq. S1.6:

$$v_{C1} = k_1 SE - (k_{-1} + k_2)C_1 = 0$$

$$\frac{(k_{-1} + k_2)C_1}{k_1} = K_{MS} C_1 = SE \qquad \text{(Eq. S2.10)}$$

Where $K_{MS} = \frac{(k_{-1}+k_2)}{k_1}$; the Michaelis-Menten constant of epiloganin. By mass balance, the free enzyme equals the initial concentration, minus the complexed enzyme: $E = E_0 - C_1 - C_2$; then:

$$K_{MS} C_1 = SE_0 - SC_1 - SC_2$$

$$(K_{MS} + S)C_1 = SE_0 - SC_2$$

$$C_1 = \frac{SE_0}{(K_{MS} + S)} - \frac{SC_2}{(K_{MS} + S)} \qquad \text{(Eq. S2.11)}$$

Now, substituting in (Eq. S2.9):

$$(v_X + v_{C2}) + v_P = k_2 C_1 = \frac{k_2 SE_0}{(K_{MS} + S)} - \frac{k_2 SC_2}{(K_{MS} + S)}$$

$$(v_X + v_{C2}) + v_P = \frac{k_2 S}{(K_{MS} + S)}(E_0 - C_2) = \frac{k_2 E_0 S}{(K_{MS} + S)}\left(1 - \frac{C_2}{E_0}\right) \qquad \text{(Eq. S2.12)}$$

Given the maximum velocity of ketologanin formation is $V_S = k_2 E_0$, and by (Eq. S2.7) $C_2 = \frac{v_P}{k_4}$ then:

$$(v_X + v_{C2}) + v_P = \frac{V_S S}{(K_{MS} + S)}\left(1 - \frac{v_P}{k_4 E_0}\right)$$

$$\frac{(v_X + v_{C2}) + v_P}{\left(1 - \frac{v_P}{k_4 E_0}\right)} = \frac{V_S S}{(K_{MS} + S)}$$

Since we can measure the maximum velocity of secoxyloganin or OME formation by assaying microsomal preparations with ketologanin, then we can obtain $k_4 E_0$ as the $V_X = k_4 E_0$, as long as the initial amount of enzyme is the same in both experiments. Then:

$$\frac{(v_X + v_{C2}) + v_P}{\left(1 - \frac{v_P}{V_X}\right)} = \frac{V_S S}{(K_{MS} + S)} \qquad \text{(Eq. S2.13)}$$

Which equals (Eq. 1) in the main text and is practically indistinguishable from (Eq. S1.7), since we cannot determine which of the measured ketologanin is complexed and which is free.

**Methods S1 Confirmation of bond stereochemistry of Oleoside Methyl Ester Standard.**

NMR spectra were measured on a 700 MHz Bruker Avance III HD (Bruker Biospin GmbH, Rheinstetten, Germany), equipped with a cryoplatform and a 1.7 mm TCI cryoprobe. MeOH-$d_3$ was used as solvent. NMR spectra were referenced to the residual solvent signals at $\delta_H$ 3.31 and $\delta_C$ 49.0. For spectrometer control and data processing Bruker TopSpin ver. 3.6.1 was used. For the selective NOESY experiment a mixing time of 600 ms was set and 4096 scans were accumulated. As transmitter frequency 1.785 ppm/1250.5 Hz was selected (methyl group adjacent to the exocyclic double bond in the oleoside methyl ester).

**Methods S2 Chemical Synthesis of Iridoid Standards.**

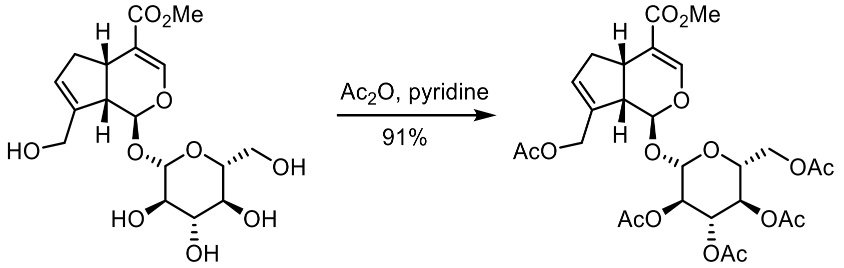


Geniposide (1g, 2.58 mmol, purchased from *Biosynth Carbosynth*.) was added to a mixed solvent of pyridine (6 mL) and acetic anhydride (3 mL), the mixed solution was stirred for 4 h at room temperature. Then the reaction mixture was concentrated in vacuum and purified by column chromatography (PE/EA = 1/1) to afford geniposide pentaacetate (1.4g, 91%). The spectroscopic data are in accordance with the literature values reported (Zhang *et al*., 2013).

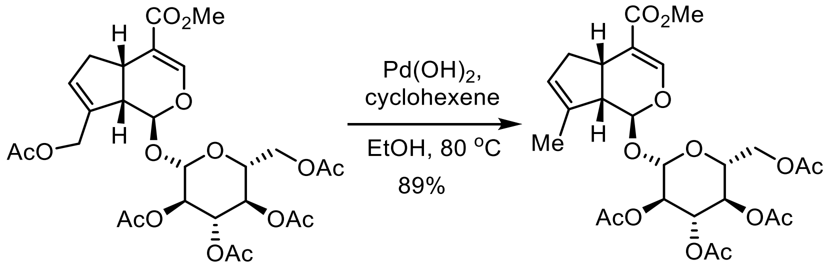


10-deoxy-geniposide
tetraacetate

Geniposide pentaacetate (900 mg) and Pd(OH)$_2$/C (90 mg) was added to a mixed solvent of cyclohexene (1.25 mL) and EtOH (2.5 mL). The mixture was refluxed for 5 h and the catalyst was filtered off. The filtrate was concentrated in vacuum and purified by column chromatography

(PE/EA = 2/1) to afford 10-deoxy-geniposide tetraacetate (720 mg, 89%). The spectroscopic data are in accordance with the literature values reported (Inoue *et al.*, 1992).

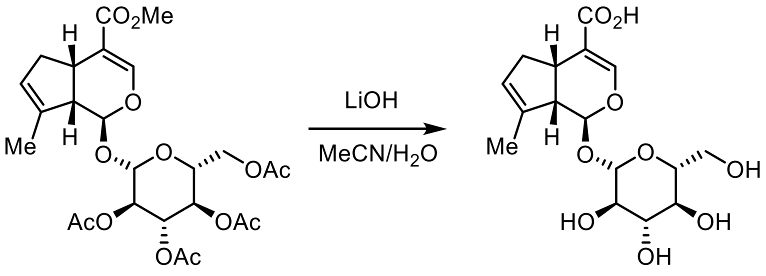


To a stirred solution of 10-deoxygeniposode tetraacetate (38 mg, 0.703 mmol) in MeCN (2.5 mL) and $H_2O$ (1 mL) was added LiOH (25 mg, 1.06 mmol) at room temperature and the resulting mixture was stirred for 3 h at 40 °C. Then the reaction was purified by column chromatography (Dowex®50WX8-200 ion-exchange resin, deionized water) directly to afford 10-deoxygeniposidic acid (27 mg, quant.). The spectroscopic data are in accordance with the literature values reported (Inoue *et al.*, 1992).

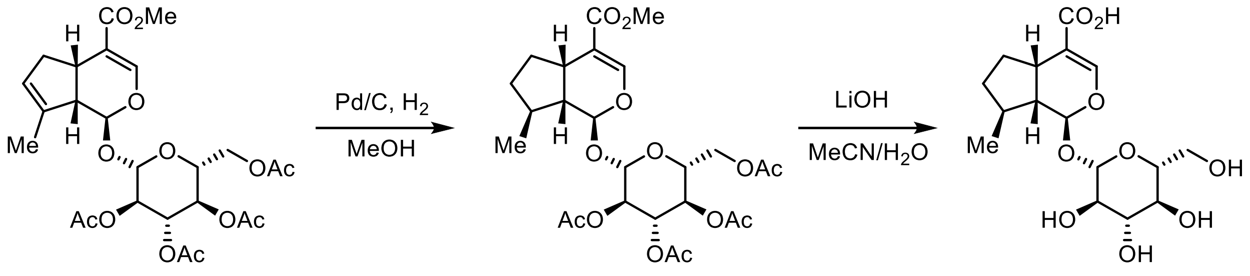


10-Deoxygeniposide (15 mg) was dissolved in MeOH (2 mL), Pd/C (1.5 mg, 5 mol%) was added and the flask was connected to a balloon filled with hydrogen and evacuated. After stirring for 1 h at room temperature under hydrogen atmosphere, the catalyst was filtered off and the solvent was evaporated. The resulted residue was used in next step directly.

To a stirred solution of above residue in MeCN (1 mL) and $H_2O$ (0.4 mL) was added LiOH (10 mg, 0.417 mmol) at room temperature and the resulting mixture was stirred for 3 h at 40 °C. Then the reaction was purified by column chromatography (Dowex®50WX8-200 ion-exchange resin, deionized water) directly to afford 7-deoxy-loganic acid (10 mg, quant.). The spectroscopic data are in accordance with the literature values reported (Inoue *et al.*, 1992).

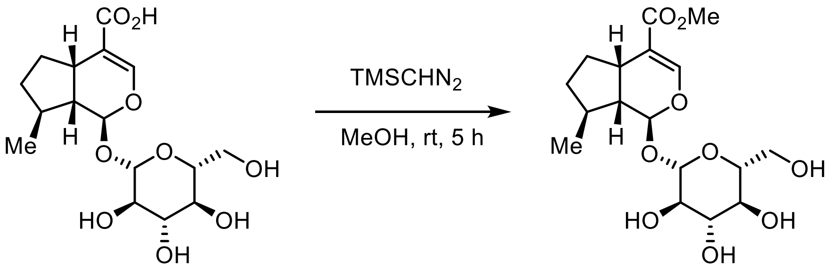


7-deoxy-loganic acid — 7-deoxy-loganin

To a stirred solution of 7-deoxy-loganic acid (2.5 mg, 0.00694 mmol) in MeOH (1 mL) was added $TMSCHN_2$ (58 uL, 0.0348 mmol, 0.6 M in hexane) at room temperature and the resulting mixture was stirred for 5 h at the same temperature. Then the reaction was purified by column chromatography (Dowex®50WX8-200 ion-exchange resin, deionized water) directly to afford 7-deoxy-loganin (2.2 mg, 85%). The spectroscopic data are in accordance with the literature values reported (Inoue *et al.*, 1992).

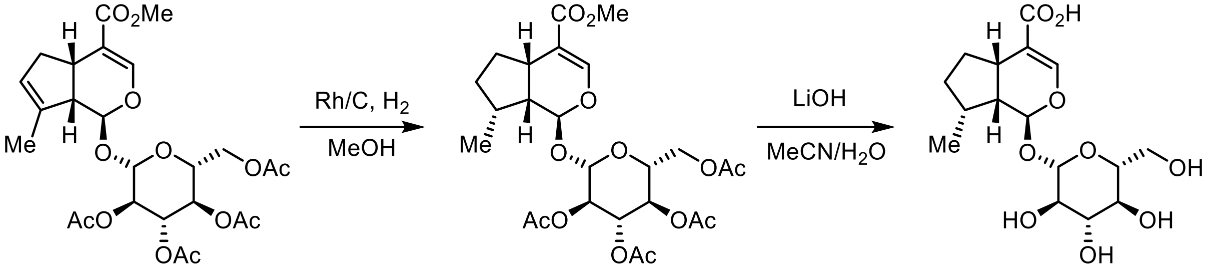


7-deoxy-8-epi-loganic acid

10-Deoxygeniposide (25 mg) was dissolved in MeOH (4 mL), Rh/C (5 mg, 5 mol%) was added and the flask was connected to a balloon filled with hydrogen and evacuated. After stirring for 2 h at room temperature under hydrogen atmosphere, the catalyst was filtered off and the solvent was evaporated. The resulted residue was used in next step directly.

To a stirred solution of above residue in MeCN (2.0 mL) and $H_2O$ (1.0 mL) was added LiOH (17 mg, 0.692 mmol) at room temperature and the resulting mixture was stirred for 3 h at 40 $^oC$. Then the reaction was purified by column chromatography (Dowex®50WX8-200 ion-exchange resin, deionized water) directly to afford 7-deoxy-8-*epi*-loganic acid (16 mg, 96%). The spectroscopic data are in accordance with the literature values reported (Nakamura *et al.*, 2000).

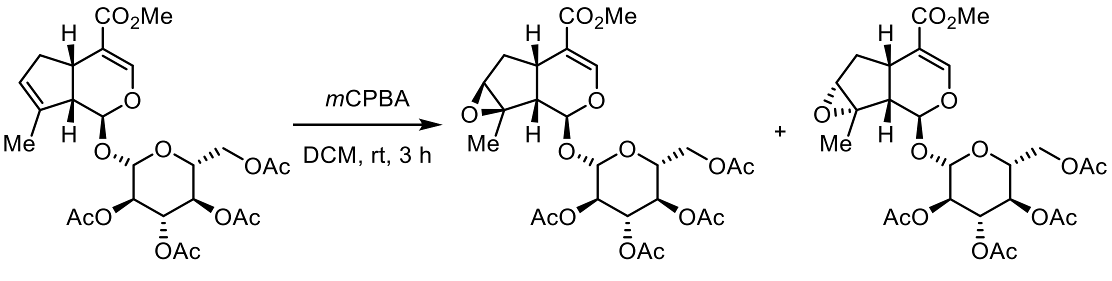


To a stirred solution of 10-deoxygeniposode tetraacetate (500 mg, 0.925 mmol) in DCM was added *m*CPBA (250 mg, 1.02 mmol) at 0 °C and the resulting mixture was stirred for 3h at room temperature. The reaction was quenched with 2N aqueous NaOH, extracted with DCM, dried over anhydrous $Na_2SO_4$ and concentrated in vacuum. The resulted residue was purified by column chromatography (PE/EA = 4/1) to afford epoxide 1 (340 mg, 66%) and epoxide 2 (110 mg, 21%). The spectroscopic data are in accordance with the literature values reported (Inouye et al., 1970).

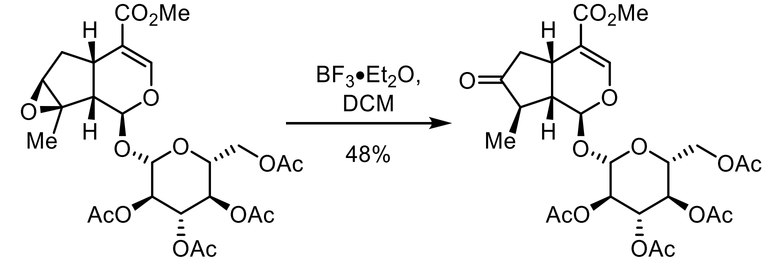


To a stirred solution of epoxide (50 mg, 0.0899 mmol) in DCM (2 mL) was added $BF_3{\cdot}Et_2O$ (0.2 mL) at room temperature and the resulting mixture was stirred for 10 min at the same temperature. The reaction was quenched with saturated $NaHCO_3$, extracted with DCM, dried over anhydrous $Na_2SO_4$ and concentrated in vacuum. The resulted residue was purified by column chromatography (PE/EA = 3/1) to afford ketone (24 mg, 48%). The spectroscopic data are in accordance with the literature values reported (Inouye et al., 1970).

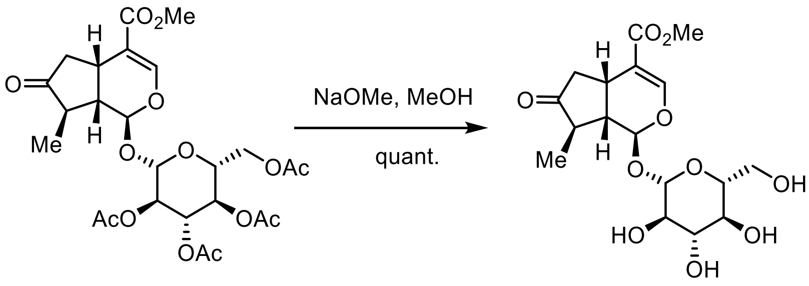

7-keto-loganin

To a stirred solution of ketone (8 mg, 0.0144 mmol) in MeOH was added NaOMe (5.4 M, 2.7 uL, 0.0144 mmol) at room temperature and the resulting mixture was stirred for 3 h at the same temperature. The reaction was purified by column chromatography (Dowex®50WX8-200 ion-exchange resin, deionized water) directly to afford 7-keto-loganin (5 mg, 90%). The spectroscopic data are in accordance with the literature values reported (Gross *et al.*, 1986).

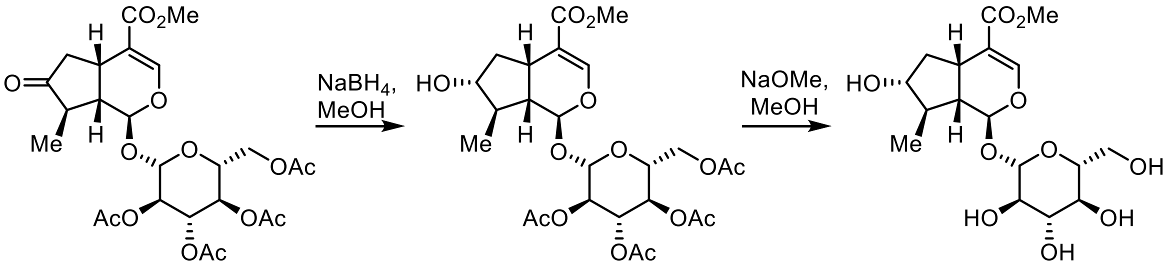

7-*epi*-loganin

To a stirred solution of ketone (7 mg, 0.0126 mmol) in MeOH (1 mL) was added $NaBH_4$ (0.5 mg, 0.0126 mmol) at room temperature and the resulting mixture was stirred for 30 min at the same temperature. The reaction was was quenched with saturated $NaHCO_3$, extracted with DCM, dried over anhydrous $Na_2SO_4$ and concentrated in vacuum. The resulted residue was used in the next step directly.

To a stirred solution of above residue in MeOH (1 mL) was added NaOMe (5.4 M, 2.3 uL, 0.0126 mmol) at room temperature and the resulting mixture was stirred for 3 h at the same temperature. Then the reaction was purified by column chromatography (Dowex®50WX8-200 ion-exchange resin, deionized water) directly to afford 7-*epi*-loganin (4.0 mg, 82%). The spectroscopic data are in accordance with the literature values reported (Itoh *et al.*, 2005).

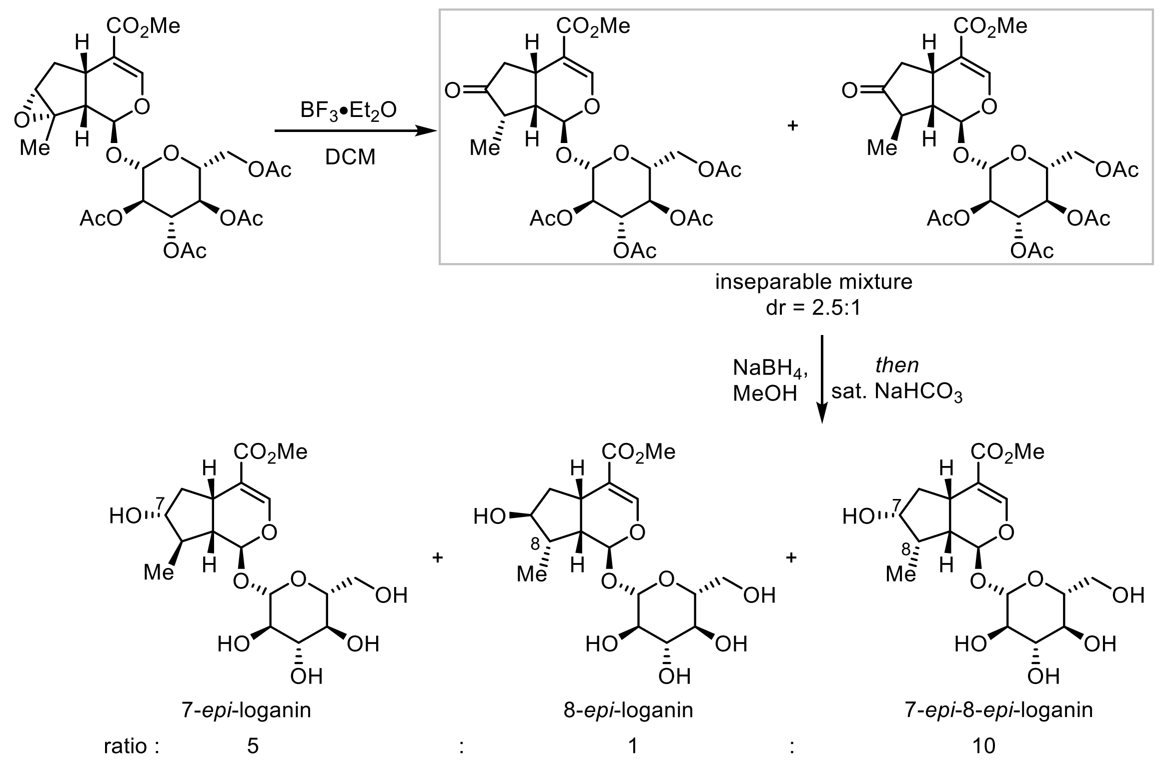


To a stirred solution of epoxide (30 mg, 0.054 mmol) in DCM (4 mL) was added $BF_3 \cdot Et_2O$ (160 uL) at room temperature and the resulting mixture was stirred for 10 min at the same temperature. The reaction was quenched with saturated $NaHCO_3$, extracted with DCM, dried over anhydrous $Na_2SO_4$ and concentrated in vacuum. The resulted residue was used in the next step directly.

To a stirred solution of above residue in MeOH (2 mL) was added was added $NaBH_4$ (2 mg, 0.054 mmol) at room temperature and the resulting mixture was stirred for 30 min at the same temperature. The reaction was was quenched with saturated $NaHCO_3$ (4 mL) and stirred overnight. Then the reaction was purified by column chromatography (Dowex®50WX8-200 ion-exchange resin, deionized water) to afford a mixture of 7-*epi*-loganin, 8-*epi*-loganin, and 7-*epi*-8-*epi*-loganin (ratio = 5 : 1 : 10, 11 mg, 52% for two steps.).

**Methods S3 Spectroscopy data for 7-epi-8-epi-loganin**

Data of 7-*epi*-8-*epi*-loganin: $^1H$ NMR (400 MHz, $CD_3OD$) δ 7.43 (s, 1H), 5.66 (d, $J$ = 5.6 Hz, 1H), 4.70 (d, $J$ = 7.9 Hz, 1H), 4.17 (dd, $J$ = 12.7, 5.7 Hz, 1H), 3.89-3.84 (m, 1H), 3.69 (s, 3H), 3.68-3.61 (m,

1H), 3.39-3.34 (m, 1H), 3.24-3.17 (m, 3H), 2.84 (dd, *J* = 18.0, 8.4 Hz, 1H), 2.61 (ddd, *J* = 14.2, 9.8, 7.4 Hz, 1H), 2.22 (td, *J* = 12.1, 6.6 Hz, 2H), 1.42-1.32 (m, 1H), 1.10 (d, *J* = 7.0 Hz, 3H);

$^{13}C$ NMR (100 MHz, $CD_3OD$) δ 168.2, 151.2, 112.2, 98.5, 95.7, 76.9, 76.6, 74.0, 73.5, 70.3, 61.5, 50.3, 42.2, 41.7, 40.5, 30.9, 9.4;
